# A Neural Mechanism for Optic Flow Parsing in Macaque Visual Cortex

**DOI:** 10.1101/2024.02.19.581050

**Authors:** Nicole E. Peltier, Akiyuki Anzai, Ruben Moreno-Bote, Gregory C. DeAngelis

## Abstract

For the brain to compute object motion in the world during self-motion, it must discount the global patterns of image motion (optic flow) caused by self-motion. Optic flow parsing is a proposed visual mechanism for computing object motion in the world, and studies in both humans and monkeys have demonstrated perceptual biases consistent with the operation of a flow parsing mechanism. However, the neural basis of flow parsing remains unknown. We demonstrate, at both the individual unit and population levels, that neural activity in macaque area MT is biased by peripheral optic flow in a manner that can at least partially account for perceptual biases induced by flow parsing. These effects cannot be explained by conventional surround suppression mechanisms or choice-related activity, and have a substantial neural latency. Together, our findings establish the first neural basis for the computation of scene-relative object motion based on flow parsing.

## Introduction

As we move through the world, our eyes are presented with a structured pattern of image motion called optic flow (Gibson, 1950; Longuet-Higgins and Prazdny, 1980). Optic flow is a rich source of self-motion information and can be used to estimate one’s heading (Britten, 2008; Van den Berg, 1992; Warren et al., 1991). However, optic flow complicates the interpretation of object motion. During self-motion, an object’s motion on the retina reflects a vector sum of image motion caused by the object’s movement in the world and optic flow due to the observer’s self-motion. To estimate object motion in the world, the brain must discount optic flow resulting from self-motion.

One mechanism for computing object motion during self-motion is known as optic flow parsing (Niehorster and Li, 2017; Rushton and Warren, 2005; Warren and Rushton, 2007; 2009a). According to the flow-parsing hypothesis, the visual system subtracts the optic flow due to self-motion such that any remaining motion represents object motion in the world (Figure 1A, B). If the visual system performs flow parsing, an observer’s perception of an object’s motion should be biased relative to its retinal motion (Warren and Rushton, 2009a), with the bias being a repulsion away from the optic flow vector at the location of the object.

**Figure 1.**
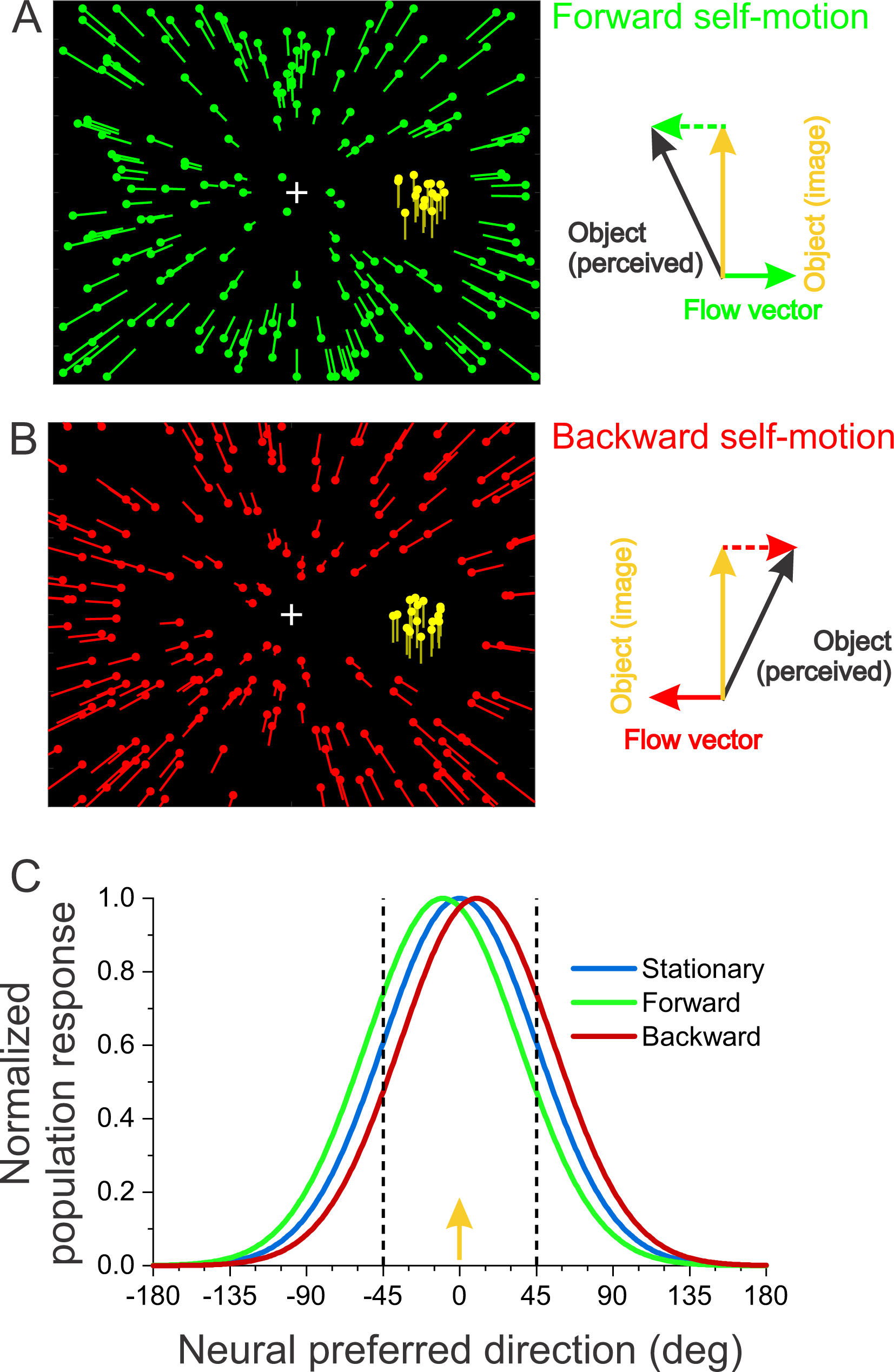
Illustration of expected perceptual biases from flow parsing and a potential neural correlate. **(A)** Schematic illustration of a stimulus condition presenting forward self-motion (green dots) and upward object motion on the screen (yellow dots). *Right*: If flow parsing occurs, the rightward flow vector at the location of the object (solid green arrow) would be subtracted, leading to a leftward bias in perceived object direction (black arrow). The yellow arrow indicates object direction in image (screen) coordinates. The dashed green arrow is the opposite of the flow vector, which is vectorially added to the image motion (yellow arrow) to obtain the expected perceived direction (black arrow). **(B)** Same as panel A except that optic flow simulates backward self-motion (red dots), leading to a rightward expected bias from flow parsing. **(C)** Hypothetical neural population response profiles in response to a presentation of an object in the right visual hemi-field moving straight upward in retinal coordinates (e.g., yellow dots in panels A and B). Each curve shows the normalized response of a population of neurons plotted as a function of each neuron’s preferred direction. When an observer is stationary (blue), the population hill of activity peaks at 0 deg (vertical motion). If the perceptual biases induced by flow parsing are reflected in this neural population response, then forward self-motion should shift the curve leftward (green) and backward self-motion should shift the curve rightward (red). As a result, a neuron that prefers a direction of -45° should have a greater response during forward self-motion than during backward self-motion. In contrast, a cell that prefers +45° would show the opposite effect.

Human psychophysical experiments have demonstrated that optic flow biases object motion perception in a manner consistent with flow parsing (Foulkes et al., 2013; Matsumiya and Ando, 2009; Niehorster and Li, 2017; Rogers et al., 2017; Warren and Rushton, 2007; 2009a; b). The direction of the induced bias is predicted by flow parsing, although the magnitude of bias is typically smaller than expected (flow-parsing gain <1, Niehorster and Li, 2017). As predicted, larger biases are observed when the object is more eccentric in the visual field (Warren and Rushton, 2009a), and the direction of the induced bias is opposite for forward and backward self-motion (Rogers *et al*., 2017). Perceptual biases also grow with self-motion speed, as expected by the flow parsing hypothesis (Niehorster and Li, 2017; Peltier et al., 2020). Flow parsing effects are seen even when optic flow is confined to the visual hemi-field opposite to that which contains the object of interest (Warren and Rushton, 2009a), suggesting a global component of the flow-parsing mechanism.

We have recently shown that motion perception of macaque monkeys demonstrates flow parsing, including all of the key features of human behavior described above (Peltier *et al*., 2020). Although biologically-plausible computational models have been proposed (Layton and Fajen, 2016b; Layton and Niehorster, 2019), the neural mechanisms underlying optic flow parsing remain unknown. Because the behavioral effects of flow-parsing are highly location-specific (Peltier *et al*., 2020), a brain area that represents the outcome of a flow-parsing mechanism is likely to contain a retinotopic motion map. The middle temporal (MT) area is a strong candidate given its retinotopic visual representation and its robust direction and speed tuning (Albright et al., 1984; Maunsell and Van Essen, 1983b; Nover et al., 2005; Van Essen et al., 1981). MT activity is correlated with perceived motion for ambiguous motion stimuli (Britten et al., 1996), plaid motion (Rodman and Albright, 1989; Stoner and Albright, 1992), illusory motion (Krekelberg et al., 2003; Luo et al., 2019), and implied motion (Kourtzi and Kanwisher, 2000; Schlack and Albright, 2007). Thus, we hypothesized that MT activity may reflect perceptual biases induced by flow parsing. Furthermore, MT has reciprocal connections with the dorsal subdivision of the medial superior temporal area (MSTd) (Maunsell and Van Essen, 1983a), an area that is highly selective for radial optic flow patterns and is causally linked to heading perception (Britten and Van Wezel, 2002; Celebrini and Newsome, 1995; Duffy and Wurtz, 1991; Gu et al., 2012; Gu et al., 2006). This connection with MSTd may allow MT to compensate for self-motion and represent object motion in the world.

If population activity in area MT accounts for the behavioral effects of flow parsing, one would expect the population activity profile to shift with the direction of self-motion simulated by optic flow (Figure 1C). This predicts that the effect of optic flow on a neuron’s response would depend systematically on the neuron’s preferred direction (dashed lines, Fig. 1C). To identify a neural mechanism for flow parsing, we recorded from small neural populations in area MT while monkeys performed a direction discrimination task in the presence of different optic flow backgrounds. We find that optic flow modulates the responses of individual units in a manner consistent with the predicted shift of the population response (Fig. 1C). Moreover, single-session population decoding shows that even small populations of MT neurons can account for a substantial portion of the behavioral effects of flow parsing. Together, our findings demonstrate a novel mechanism for computing scene-relative object motion based on flow parsing.

## Methods

### Subjects and surgery

Two male rhesus monkeys (*Macaca mulatta*) participated in the experiment. A head restraint device was implanted according to standard aseptic surgical procedures under gas anesthesia. A Delrin (Dupont) ring was attached to the skull with dental acrylic cement and anchored with bone screws and titanium inverted T-bolts (see Gu *et al*., 2006 for details). To monitor eye movements, a scleral search coil was implanted under the conjunctiva of one eye.

To guide electrodes to area MT, a Delrin recording grid was fastened inside the head-restraint ring using dental acrylic. The recording grid (2 × 4 × 0.5 cm) contained a dense array of holes spaced 0.8 mm apart. Under anesthesia and using sterile technique, small burr holes (∼0.5 mm diameter) were drilled vertically through the recording grid to allow electrodes to penetrate the brain through transdural guide tubes. All surgical procedures and experimental protocols were approved by the University Committee on Animal Resources at the University of Rochester.

### Experimental apparatus

Monkeys were seated in custom-built primate chairs with their heads restrained. The chair was fastened onto a six degree-of-freedom motion platform (MOOG 6DOF2000E); however, in these experiments, the platform remained stationary. A field coil frame (C-N-C Engineering) was mounted to the top of the motion platform to monitor eye movements using the scleral search coil technique.

Visual stimuli were rear-projected onto a 60 × 60 cm tangent screen using a projector (Christie Digital Mirage S+3K) that was mounted on the motion platform. The screen was affixed to the front of the field coil frame, roughly 30 cm in front of the monkey (monkey M: 31.7 cm from eyes to screen; monkey P: 33.0 cm from eyes to screen). As a result, the screen subtended approximately 90 × 90° of visual angle. To restrict the monkey’s field of view to visual stimuli on the screen, the sides and top of the field coil frame were covered with black matte material.

### Electrophysiological recordings

#### Electrode positioning system

Extracellular neural activity was recorded using V-Probe multi-site linear electrode arrays (Plexon). The probes had 24 channels with 50 µm spacing between channels. The position of the probe was controlled using the EPS electrode positioning system and the Flex MT microdrive mounting system (Alpha Omega). The Flex MT mounting ring was secured to the monkey’s head restraint, and a microdrive tower was mounted onto this ring. This tower held the V-Probe and guide tube, while connecting to the EPS system to drive the electrode array. The sterilized V-Probe was front-loaded into a transdural guide tube and then secured to the microdrive tower. The tower was then affixed to the mounting ring in a position that aligned the guide tube vertically with the appropriate grid hole. The guide tube was then lowered manually through the grid hole until the resistance of the dura mater was felt. The entire tower, including the guide tube and electrode array, was then lowered ∼2-3 mm, allowing the guide tube to puncture the dura mater.

#### Neural signal processing system

Neural signals were amplified and bandpass filtered (350 Hz – 3446 Hz, Blackrock Microsystems). Spike waveforms and raster plots for all channels were monitored online. Spike detection thresholds were set manually for each channel to capture multi-unit activity with a spontaneous firing rate of ∼50-100 spikes/second. The raw voltage signals from the probe were digitized and stored to disk at 30 kHz for offline analysis. Because spike thresholding was performed again offline before spike sorting and analysis, the manual spike thresholds were set online simply to map receptive fields and to tailor stimuli to neuronal preferences.

Neural signals were analyzed offline using Plexon Offline Sorter to determine a spike detection threshold and to perform spike sorting. Waveforms corresponding to candidate neural events were detected when the raw voltage trace reached local minima at least 3 standard deviations below the mean of the signal for each channel. Waveform snippets were extracted as 48 samples over 1.6 ms and aligned such that 16 samples were taken before the threshold was reached. These waveforms were then sorted in a 2-dimensional feature space using the built-in t-distribution expectation-maximization scanning method. As there was a tendency of the Plexon Offline Sorter to overestimate the number of unique units, the sorting of each channel was reviewed manually in a 2-dimensional feature space and units were merged as necessary. Each channel yielded one multi-unit with possibly one or more single units. The vast majority of recordings were multi-units; note, however, that multiunit signals in area MT typically show robust tuning properties that are closely matched with single units at the same location (DeAngelis and Newsome, 1999)

#### Identifying the location of area MT

The location of area MT was initially estimated from structural MRI scans and a standard macaque atlas (Van Essen et al., 2001). Area MT was identified as a region in the posterior bank of the superior temporal sulcus (STS), typically centered ∼16 mm lateral to the midline and ∼3 mm posterior to the interaural plane. Before using a V-Probe, tungsten microelectrodes (FHC) were used to identify MT by mapping the response properties in locations corresponding to the region identified by MRI. Electrode penetrations were informed by the pattern of activity as the electrode passed through gray matter and white matter, as well as the response properties of neurons to visual stimuli. As the electrode approached the STS, it typically encountered neurons with large receptive fields that were selective for direction of visual motion, characteristic of the dorsal division of the medial superior temporal (MSTd) area (Duffy and Wurtz, 1991). This activity was typically followed by a very quiet region, indicative of the lumen of the STS, and then area MT as the next region of gray matter. Receptive fields in MT were markedly smaller than those in MSTd, and their sizes scaled approximately linearly with receptive field eccentricity (Albright and Desimone, 1987). Across MT, neurons typically exhibited tuning to direction, speed, disparity, and receptive field location that changed gradually with the depth of the electrode, consistent with the previously documented organization of MT (Albright *et al*., 1984; DeAngelis and Newsome, 1999).

In recording experiments, the V-Probe was lowered until activity characteristic of MT was centered on the probe’s channels, after which the probe was left to settle for 1-1.5 hours. As the brain settled around the probe, the MT activity sometimes shifted toward shallower channels. This tendency was combatted by retracting the probe 10-50 µm at a time to keep MT centered on the channels.

### Visual stimuli

Visual stimuli simulated the motion of an independently moving object during forward or backward self-motion. Stimuli were generated by software written in Visual C++, using the OpenGL 3D graphics rendering library. An OpenGL camera located at the same position as each of the animal’s eyes generated the planar image projection shown to each eye. To simulate depth in the stimulus, the stimulus was rendered stereoscopically as a red/green anaglyph, and the animal viewed it through red and green filters (Kodak Wratten2 #29 and #61, respectively). Visual stimuli lasted for 2 s in all recording sessions except for the first three sessions in one monkey; in these experiments, the stimulus duration was 1.2 s.

#### Object motion

Object motion was represented by random dots moving coherently within a circular aperture positioned approximately in the middle of the receptive fields of the recorded MT units, as determined with the receptive field mapping protocol described below. This patch of random dots (hereafter typically referred to as the “object”) was rendered at the same distance as the fixation point (i.e., centered within the plane of the screen), and its size was determined to be the approximate size of the receptive fields of several channels sampled online. The object was a nearly flat disk, with a front-to-back simulated visual depth of 0.1 cm. Dots within the object were triangles 0.15 cm wide and 0.15 cm tall, and they were distributed with a density of 20 dots/cm^3^.

Object motion within a stationary aperture was used so that the object stayed on the receptive field throughout the trial, reducing response modulations due to varying luminance within the receptive field or stimulation of the inhibitory surround. The stationary aperture ensured that any observed difference in firing rate between object motion directions was due to motion within the aperture and not to changes in position of the object’s boundaries. We also measured behavior in some sessions using moving objects in which the object boundary translated, and we observed no qualitative differences in biases or discrimination thresholds induced by optic flow. Thus, flow parsing appears to function similarly for “objects” with either stationary or moving boundaries.

The object moved in the fronto-parallel plane in one of 11 directions centered around straight upward. For monkey M, object directions ranged from -40° to +40° around straight upward, spaced linearly in steps of 8°. For monkey P, object directions ranged from -20° to +20°, spaced in steps of 4°. Linear spacing of object directions was used to allow for equal resolution in measuring perceptual biases within the range of tested directions. The 11 object directions were interleaved randomly within a single block of trials. In conditions with simulated self-motion, the object moved in depth with the OpenGL camera, staying at the same depth relative to the moving camera, and thus keeping the location and size of the object fixed on the retina. The direction and speed of object motion were therefore defined in screen coordinates. Since the fixation target remained fixed on the screen during simulated self-motion, any given direction of object motion was identical on the retina (assuming perfect fixation) for the different optic flow conditions. Thus, any observed differences in firing rate between optic flow conditions could be attributed to the surrounding pattern of optic flow.

Dots within the object moved coherently, following a Gaussian velocity profile with a standard deviation equal to 1/6^th^ of the duration of the visual stimulus (σ = 0.2 s for 1.2-s trials, 0.33 s for 2-s trials), hitting the peak speed around the middle of the trial. Generally, this velocity profile was scaled such that the peak speed was 10°/s, selected because it is approximately the median preferred speed of MT neurons (Nover *et al*., 2005). However, in sessions where there were channels on the V-Probe that did not respond at all to slow speeds, the peak speed was increased to 16°/s or 20°/s.

#### Self-motion

A three-dimensional cloud of background dots surrounded the object, extending in depth from 5 cm to 55 cm from the eyes. Patterned motion of these background dots was used to visually simulate self-motion (see Movie #1). Each simulated self-motion was a pre-programmed straight trajectory (forward or backward) that was not under control of the subject. During simulated self-motion, the depth range of the dots (relative to the OpenGL camera) remained fixed through the use of near and far clipping planes in OpenGL. The background dots were triangles with height and width of 0.1 cm, distributed with a density of 0.002 dots/cm^3^. In the first four sessions with monkey P, the background dots were points 3 × 3 pixels in size, and their retinal projections did not vary in size according to their distance from the monkey. In subsequent sessions, these fixed-size dots were replaced with triangles whose image sizes varied inversely with their distance from the monkey. This added a monocular depth cue to the optic flow, simulating self-motion more realistically.

In most trials, background motion was generated by placing an OpenGL camera at the location of the monkey’s eye and moving the camera along the trajectory of the simulated self-motion. In the *forward* self-motion condition, the background dots expanded radially from the central fixation point, simulating forward self-motion. In the *backward* condition, the background dots contracted toward the fixation point, simulating backward self-motion. In a set of control trials (*stationary* condition), the background dots were static for the duration of the trial to indicate no self-motion. These three self-motion conditions were crossed with the 11 object directions, and all conditions were randomly interleaved within a single block of trials.

Self-motion speed followed a Gaussian velocity profile with a standard deviation equal to 1/6 of the trial duration (σ = 0.2 s for 1.2-s trials, 0.33 s for 2-s trials), hitting the peak speed around the middle of the trial. This velocity profile was identical in shape and timing to the object’s velocity profile; therefore, object speed and simulated self-motion speed are proportional throughout the trial.

Self-motion speed, along with the object’s horizontal location and speed, determines the image velocity of the flow vector at the location of the object (Longuet-Higgins and Prazdny, 1980), and thus the predicted perceptual bias due to flow-parsing (Peltier *et al*., 2020). In our experiments, the speed of self-motion was chosen for a given object location and speed to keep the predicted perceptual bias at a specified value. Self-motion was, therefore, faster in sessions with objects that were less horizontally eccentric and in sessions with faster object motion. The predicted bias was specified for each session, and it was set to be either ±10° or ±15° so that the total predicted shift between forward and backward self-motion would be either 20° or 30° (assuming perfect flow parsing with a gain of unity). For monkey M, the predicted bias was ±10° in all experiments. Because monkey P generally exhibited smaller biases that decreased toward 0 over an extended period of training (Suppl Figure 1B), faster self-motion was used in later experiments to elicit a robust perceptual bias. For monkey P, the predicted bias was ±10° in early experiments and ±15° in several later experiments.

A circular mask was placed around the object to block out background dots directly surrounding the object. The purpose of the mask was to prevent the monkey from making judgments of object motion based solely on local motion comparisons between the object and the immediately surrounding optic flow. The background mask also limited the effect of center-surround interactions on MT responses. The size of the mask was determined by a mask ratio parameter, which is the ratio of the mask’s diameter to the object’s diameter. In these experiments, the mask ratio stayed constant within each session and ranged between 2 and 3 across sessions.

### Experimental protocol

#### Preliminary measurements

##### Manual mapping of MT

Once the V-Probe had settled in area MT, the receptive field and tuning properties of several channels spanning the length of the probe were examined individually using manually controlled patches of random dots. The parameters of the patch were manipulated to determine approximate receptive field location and size, as well as estimates of preferences for direction, speed, and disparity.

##### Tuning protocols

Quantitative measurements were then taken of direction tuning, speed tuning, and receptive field location and size. While the activity from all channels was saved for offline analysis, the activity of four channels distributed along the length of the probe was monitored online during these tuning measurements. The tuning of these four channels was used to inform stimulus parameter selection in subsequent tuning protocols.

Direction tuning was measured with patches of random dots that drifted coherently in one of 8 directions separated by 45°, in order to determine the direction preferences of all neurons. The estimated preferred direction (determined by eye from the online tuning curve) was set to be the stimulus direction in subsequent tuning measurements. When preferred directions varied substantially across channels in a recording session, an intermediate direction (chosen to activate the most units) was selected as the preferred direction. If preferred directions differed so much that an intermediate direction would not elicit responses from any channels, subsequent tuning measurements were taken more than once, testing responses to two different directions that together could drive responses from most, if not all, channels.

Speed tuning was measured with patches of random dots that moved coherently in the preferred direction at 0, 0.5, 1, 2, 4, 8, 16, and 32°/s. The preferred speed was estimated manually from tuning curves plotted online from neural activity recorded on the four selected channels, and it was used as the stimulus speed in subsequent tuning measurements. If there appeared to be substantial variation in speed preferences, an intermediate speed was used. It never occurred that a single speed could not be found that would elicit a response from all four of the channels monitored online.

The spatial profile of the receptive field was measured with patches of random dots presented at locations on a 4 × 4 grid. The grid was centered on the manually estimated receptive field center and covered an area twice as wide and as long as the manually estimated diameter of the receptive field. Responses were fitted with a 2-dimensional Gaussian function to estimate the center and size of the receptive field. The mean of the receptive field centers of the four channels that could be viewed online determined the location of the object during the size tuning measurement and the flow-parsing experiment.

Size tuning was measured with patches of random dots that moved in the preferred direction and speed with patch diameters of 2, 4, 8, 16, 32, and 64° of visual angle. The approximate mean optimal stimulus size among the four channels viewed online determined the object’s size in the flow-parsing experiment. Optimal sizes varied from 8° to 32° in diameter depending on stimulus eccentricity, with a median diameter of 20°.

#### Flow-parsing task

The animal’s task was to judge whether a patch of dots (the “object”) moved rightward or leftward, relative to vertical, in the presence of optic flow simulating forward or backward self-motion (see Suppl. Figure 2 and Fig. 2 of (Peltier *et al*., 2020) for more details). At the beginning of each trial, a fixation target appeared in the center of a blank screen. The monkey had to maintain fixation within a 2.5-2.8° (full width) box surrounding the fixation target for the duration of the trial. 215 ms after fixation was achieved, the visual stimulus appeared. The object and background dots appeared simultaneously, started to move following the Gaussian velocity profile described above, and then disappeared when the object motion and self-motion concluded. At this time, the fixation point disappeared and two choice targets appeared 10° to the left and right of center. The monkeys indicated whether they perceived leftward or rightward object motion (relative to the vertical reference) with a saccadic eye movement to one of the two choice targets. Correct responses, based on the direction of object motion in screen coordinates, were followed by a liquid reward (0.2-0.4 ml).

**Figure 2.**
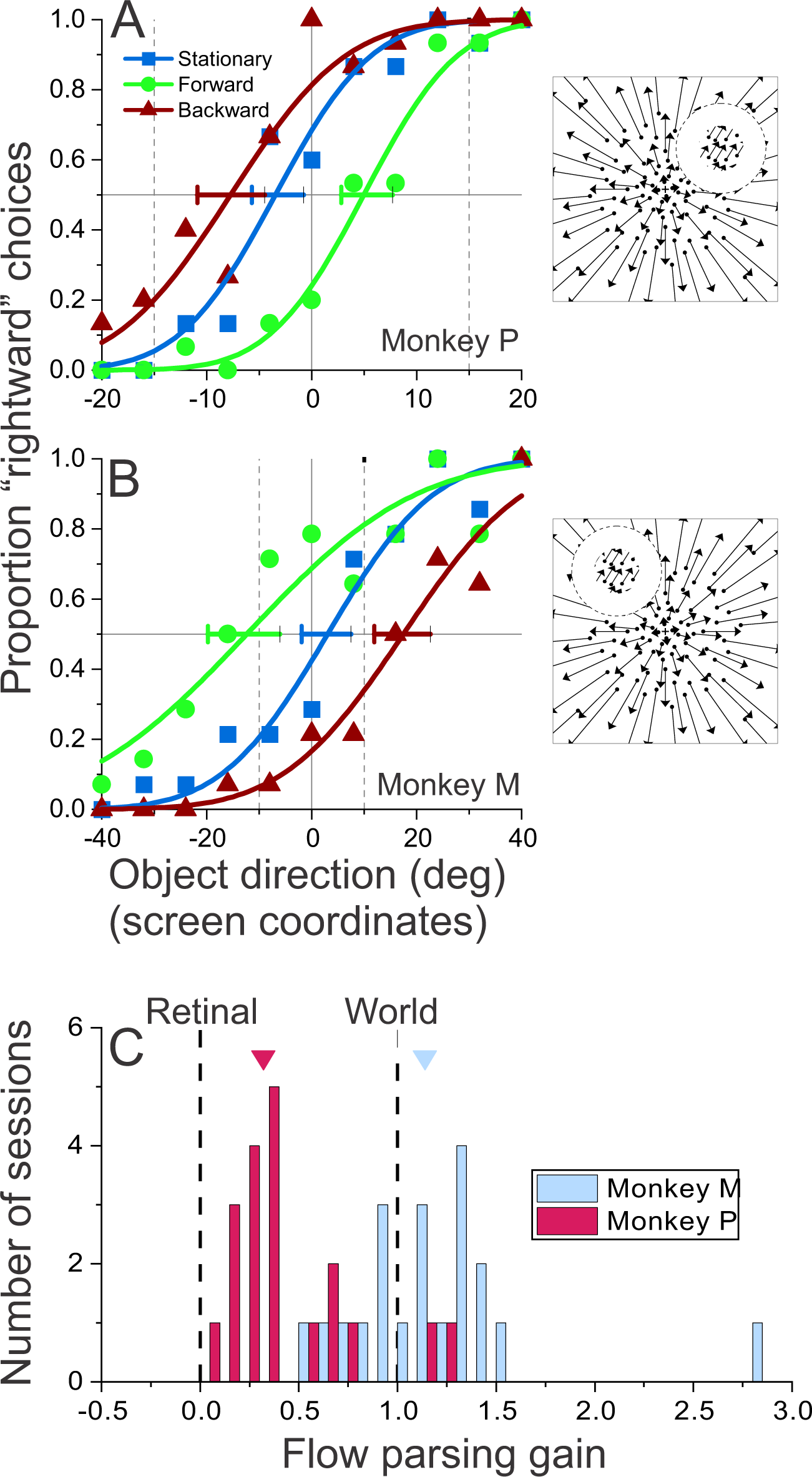
Optic flow systematically biases object motion perception in monkeys. **(A)** Psychometric functions from a recording session in which monkey P discriminated object direction in the presence of optic flow. Symbol shape and color denote data from the stationary (blue squares), forward (green circles), and backward (red triangles) self-motion conditions. Smooth curves show fits of a cumulative Gaussian function to the data points. Horizontal error bars indicate 95% confidence intervals on the PSEs, and the dashed vertical lines indicate the expected PSEs for complete flow-parsing (FP gain = 1). Because the object was presented in the right visual hemi-field, monkey P’s perception of object motion was biased leftward during forward self-motion and rightward during backward self-motion. **(B)** Data from a recording session in which monkey M discriminated object motion in the left hemi-field, leading to an opposite pattern of perceptual biases. **(C)** Distributions of flow-parsing gains (observed/expected PSE shift) for 20 recording sessions from monkey M (teal) and 19 recording sessions from monkey P (purple). Downward-pointing triangles indicate the median flow-parsing gains for each animal.

The presence of optic flow was expected to bias the monkey’s reports of object direction if it was performing a flow-parsing operation (Peltier *et al*., 2020). As a result, the monkey’s report of object motion direction in the stationary condition could flip from rightward to leftward or vice versa during self-motion. This reversal could occur in conditions for which the object’s motion in screen coordinates is rightward while the object’s motion in world coordinates is leftward, or vice versa (see (Peltier *et al*., 2020) for more details). This corresponds to conditions in which the horizontal component of the optic flow vector implied at the location of the object is in the same direction and greater in magnitude than the horizontal component of object motion (assuming a flow parsing gain of unity). To avoid reinforcing perceptual reports in one coordinate frame over the other, subjects received a reward randomly on 70% of these trials (see Figure 12 of (Peltier *et al*., 2020)). On all other trials, in which object directions were consistent in screen and world coordinates, subjects were rewarded on 95% of correct trials.

### Data Analyses

#### Behavioral analyses

For each experimental session, a psychometric function was computed for each optic flow condition to represent the proportion of rightward choices as a function of object direction. The probability of a rightward choice given the object direction was calculated as a cumulative Gaussian distribution, given by:

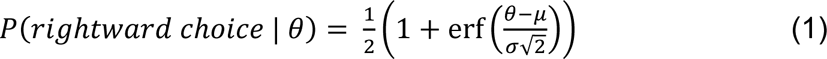

where *θ* is the object’s direction of motion, *µ* is the mean of the Gaussian distribution, *σ* is the standard deviation of the distribution, and erf(*x*) is the Gauss error function given by:

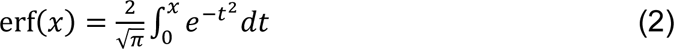

Parameters *µ* and *σ* were optimized to minimize the sum squared error between the predicted proportion of rightward choices and the recorded proportion of rightward choices. To calculate confidence intervals around each *µ*, 200 bootstrapped samples of the behavioral data were computed. Each sample was fitted with a cumulative Gaussian function, and a 95% confidence interval was computed using percentiles of the bootstrap distribution. Incorporating a lapse rate into fits of the psychometric function was not found to improve the fits and was thus not included in the main analysis.

The mean of the Gaussian distribution, *µ*, represents the object motion on the screen at which the monkey makes 50% rightward choices and 50% leftward choices, also known as the point of subjective equality (PSE). The effect of optic flow on perceived object motion was measured as the shift in PSE between forward and backward optic flow conditions, given by:

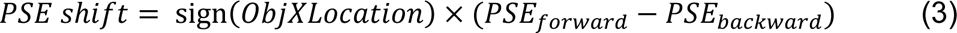

where sign(*ObjXLocation*) is +1 for objects located in the right visual field and -1 for objects located in the left visual field, and *PSE_forward_* and *PSE_backward_* are the PSEs of the psychometric function for the forward and backward optic flow conditions, respectively. The expected effect of flow parsing depends on the direction of optic flow vectors that would have been at the location of the object had they not been masked, which depends on the object’s location in the visual field. Thus, the sign of the PSE shift depends on the object’s horizontal location such that it is positive if it is in the direction predicted by flow parsing. Confidence intervals on PSE shifts were computed using the same bootstrapped samples used to compute PSE confidence intervals. For each of the 200 samples, the PSE shift was computed, and a 95% confidence interval was computed using percentiles of the bootstrap distribution.

PSE shifts were compared to the shifts that are predicted by flow-parsing through the computation of a flow-parsing gain (FP gain), given by

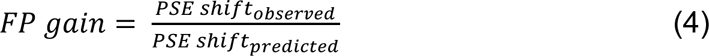

where *PSEshift_observed_* denotes the measured PSE shift (Equation 3) and *PSEshift_predicted_* denotes the PSE shift predicted by perfect flow parsing. The computation of predicted PSE shift is described in detail elsewhere (Peltier *et al*., 2020). Flow-parsing gain will be 0 if optic flow does not produce any perceptual biases, meaning that the subject is reporting object motion in retinal coordinates. Flow-parsing gain will be 1 if the biases induced by optic flow match those that are predicted by the flow-parsing hypothesis, indicating that the subject is fully compensating for self-motion and reporting object motion in world coordinates.

#### Neural analyses

##### Unit inclusion criteria

*Exclusion of units without direction tuning*: Units were included in analysis only if they exhibited statistically significant direction tuning. For each unit, firing rates collected in the direction tuning protocol were compared using a Kruskal-Wallis one-way analysis of variance to test whether the distributions of responses elicited by each direction all came from the same distribution. If the distributions of firing rates did not differ significantly between directions (*p* > 0.05), the unit was determined not to have significant direction tuning and it was excluded from subsequent analyses.

*Exclusion of units without structured receptive fields*: Firing rates recorded during the receptive field (RF) mapping protocol were fitted with a 2-dimensional (2D) Gaussian function. A unit’s firing rate at location (*x, y*) is modeled as:

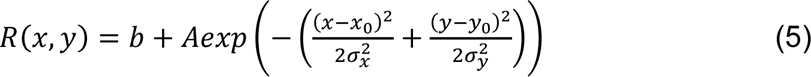

where *b* is the spontaneous firing rate, *A* is the response amplitude, *x_0_* is the horizontal location of the RF center, σ_x_ is the horizontal extent, *y_0_* is the vertical location of the RF center, and σ_y_ is the vertical extent. Parameters were optimized to minimize the sum squared error between measured and predicted firing rates. If a Spearman’s rank correlation between measured and the predicted firing rates failed to reach significance (permutation test, *p* < 0.05), the unit was excluded from further analysis.

Because the stimuli used to map MT receptive fields were approximately half the diameter of the receptive field, RF sizes are generally overestimated using this RF mapping protocol. To account for this spatial blurring effect, deconvolution of the fitted receptive fields was performed to compute a more accurate receptive field size. The 2-dimensional Gaussian fits were extrapolated onto a spatial domain that was double the length and width of the RF mapping grid, using Equation 5. Then, the horizontal and vertical components of the receptive field were deconvolved separately. For the horizontal component, a cross-section was taken at the peak vertical value. This 1-dimensional Gaussian function was then deconvolved with a boxcar function having the width of the RF mapping stimulus. The result was then fitted with a Gaussian function, given by:

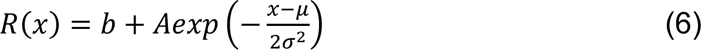

where *b* is the baseline, *A* is the amplitude, *µ* is the center location of the RF, and *σ* is the standard deviation of the Gaussian RF profile. The σ parameter from this fit was used as the adjusted *σ_x_* for the 2-dimensional receptive field. Similarly, an adjusted *σ_y_* was computed by deconvolving a cross-section taken at the peak horizontal value and fitting the output with a Gaussian function. Deconvolution typically reduced *σ* values by about 25%.

*Exclusion of units based on receptive field overlap with the optic flow background*: The purpose of the background mask was to prevent units from responding directly to the background optic flow, so units were excluded from analysis if their receptive fields had substantial overlap with the background optic flow field. We quantified the amount of overlap between receptive fields and the background mask with a metric called Receptive Field Inside Mask (RFIM). To capture the entire receptive field, we used the fitted parameters from Equation 5 to compute the receptive field over a region 3 times as wide and as tall as the range of locations tested in the RF mapping protocol. The receptive field was computed with 0.5° resolution, using parameters *b*, *A*, *x_0_*, and *y_0_* from the pre-deconvolved fit along with *σ_x_* and *σ_y_* from the deconvolved fit. The minimum value of the extended receptive field was subtracted from all points to make the minimum of the receptive field zero.

The location of the background mask was represented as a bit mask of the same size and resolution as the modeled receptive field, with values of 1 representing points within the mask and values of 0 representing points overlapping with optic flow. The overlap between the receptive field and background mask was computed as the pointwise product of the fitted receptive field and the bit mask. The products were then summed over all points and normalized by the entire area of the receptive field. This computation of RFIM is given by:

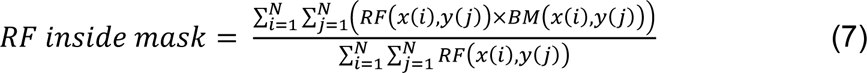

where *RF(x(i), y(j))* is the value of the unit’s receptive field at point *(x(i), y(j))* minus the minimum value and *BM(x(i), y(j))* is the value of the bit mask at the same point.

This computation of RFIM assigns a higher weight to locations that elicit higher firing rates, representing the center of the receptive field, and less weight to locations at the edge or outside the receptive field. Units were excluded from analysis if RFIM was less than 0.75, indicating that less than 75% of the response density fit within the background mask. Varying this criterion for RFIM from 0.6 to 0.9 had little impact on decoding results.

##### Spike counting window

Unless specified otherwise, firing rates were calculated by summing spike counts within a 1000-ms window centered on the peak of the population response. The peak population response occurred approximately 1200 ms after the stimulus onset, 110 ms after the peak object speed was reached. The window was therefore determined to be 700 ms to 1700 ms after stimulus onset, thus capturing most of the visually driven responses while excluding transient responses to the onset of the stimulus (see Figure 5A). As described below, a different analysis window was utilized for the computation of Flow Modulation Index and decoding analyses, based on the observed time course of FMI (Figure 5C).

##### Surround suppression index (SSI)

Firing rates recorded during the size tuning protocol were fitted with two tuning curves. The first was a single error function (DeAngelis and Newsome, 1999; DeAngelis and Uka, 2003) representing a unit’s response to stimulus diameter, *w*, given by:

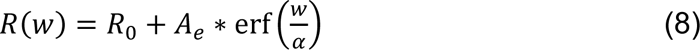

where *R_0_* is the baseline response, *A_e_* is the excitation amplitude, *α* affects the slope of the tuning curve, and erf(*x*) represents the Gauss error function (Equation 2). Equation 8 best represents the size tuning of a unit without surround suppression, as there is no peak representing a preferred stimulus size.

The second function used to fit size tuning curves was a difference-of-error (DoE) functions (DeAngelis and Newsome, 1999; DeAngelis and Uka, 2003), given by:

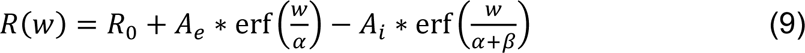

Here, parameters *R_0_*, *A_e_*, and *α*, as well as the error function erf(*x*), are as defined in Equation 8. *A_i_* represents the amplitude of inhibition, and *β* affects the slope of inhibition. The DoE function best characterizes the size tuning of units with surround suppression, as they produce a peak response for an intermediate stimulus size, with decreasing responses for larger stimuli.

The errors of the fits from Equations 8 and 9 were compared using a sequential F test, and a unit was determined to have significant surround suppression if the DoE function yielded a significantly better fit than the single error function (*p* < 0.05). If the DoE function produced a better fit, the unit’s optimal stimulus size was determined as the peak of the DoE fit. If there was no significant surround suppression, the unit’s optimal stimulus size was taken to be the size at which the single error function fit reached 90% of its maximal value (DeAngelis and Uka, 2003). A surround suppression index (SSI) was calculated for each unit as follows:

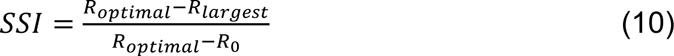

where *R_optimal_* denotes the unit’s response to its optimal stimulus size, *R_largest_* represents its response to the largest presented stimulus, and *R_0_* is the unit’s baseline response without any stimulus. SSI will be close to 1 for units that exhibit strong surround suppression, and it will be 0 for units that do not exhibit any surround suppression. Units that did not exhibit significant surround suppression, based on the sequential F-test, were assigned an SSI of 0.

##### Horizontal direction discrimination index (HDDI)

The selectivity of MT units for direction during the discrimination task (i.e., a preference for rightward or leftward object motion) was computed as a horizontal-direction discrimination index (HDDI), calculated as follows (adapted from DeAngelis and Uka, 2003; Prince et al., 2002):

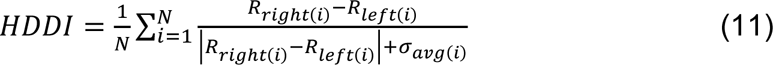

For each pair of the *N* = 5 object motion directions symmetric around 0 (e.g. ±4 degrees), the difference in mean firing rate between rightward (*R_right_*) and leftward (*R_left_*) object directions was calculated among trials in which there was no self-motion. This difference was then normalized by the sum of the magnitude of the difference and the average standard deviation of firing rates between rightward and leftward object directions (*σ_avg_*), also calculated from stationary background trials. This value was averaged over the 5 pairs of directions to determine the HDDI. HDDI ranges from -1 to 1, with negative values assigned to units that respond more strongly to leftward motion and positive values assigned to units that respond more strongly to rightward motion.

##### Flow-modulation index (FMI)

The effect of optic flow on a unit’s response was computed as a flow modulation index (FMI), given by:

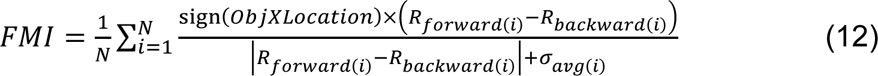

For each of the *N* = 11 object directions used in the experiment, we calculated the difference in mean firing rate between forward (*R_forward_*) and backward (*R_backward_*) self-motion conditions. This difference was normalized by the magnitude of the difference added to the average standard deviation in firing rates between forward and backward self-motion conditions. This normalized difference was averaged over the 11 object directions to compute the FMI. FMI ranges from -1 to 1, with the magnitude of the value indicating the strength of the effect of optic flow on firing rates. As with the computation of PSE shifts, the horizontal position of the object (*ObjXLocation*) is incorporated into calculation of the FMI because the expected effect of optic flow on neural responses depends on the direction of optic flow surrounding the object, which depends on the object’s location.

Because FMI is computed as an average across the different object directions, it will only tend to be substantially different from zero if there is a fairly consistent difference in response between forward and backward optic flow conditions across the different object directions. However, because the range of object directions used in the discrimination task is restricted (to ±20° for monkey P and ±40° for monkey M), we expect a consistent difference in response between the two self-motion directions for neurons with direction preferences that are substantially away from the vertical task reference (e.g., dashed vertical lines in Figure 1C).

After observing that FMI developed more slowly than HDDI (Figure 5B,C), we determined a new time window to compute FMI that was shifted toward later times than other spike rate analyses. We calculated a time course of FMI for each unit by splitting trials into 50-ms windows and computing FMI within each window. Because the strength of FMI effects is related to HDDI (Figure 3D), we separated units into two groups based on whether HDDI was greater than or less than 0, and we computed an average FMI time course for each group. The difference in time course between groups was calculated, and the absolute value of that difference was computed. We determined the time window with the strongest FMI information as the full width of the difference time course at half maximum. This window was 850-1150 ms after stimulus onset for sessions with 1200-ms trials and 1250-1750 ms after stimulus onset for sessions with 2000-ms trials.

**Figure 3.**
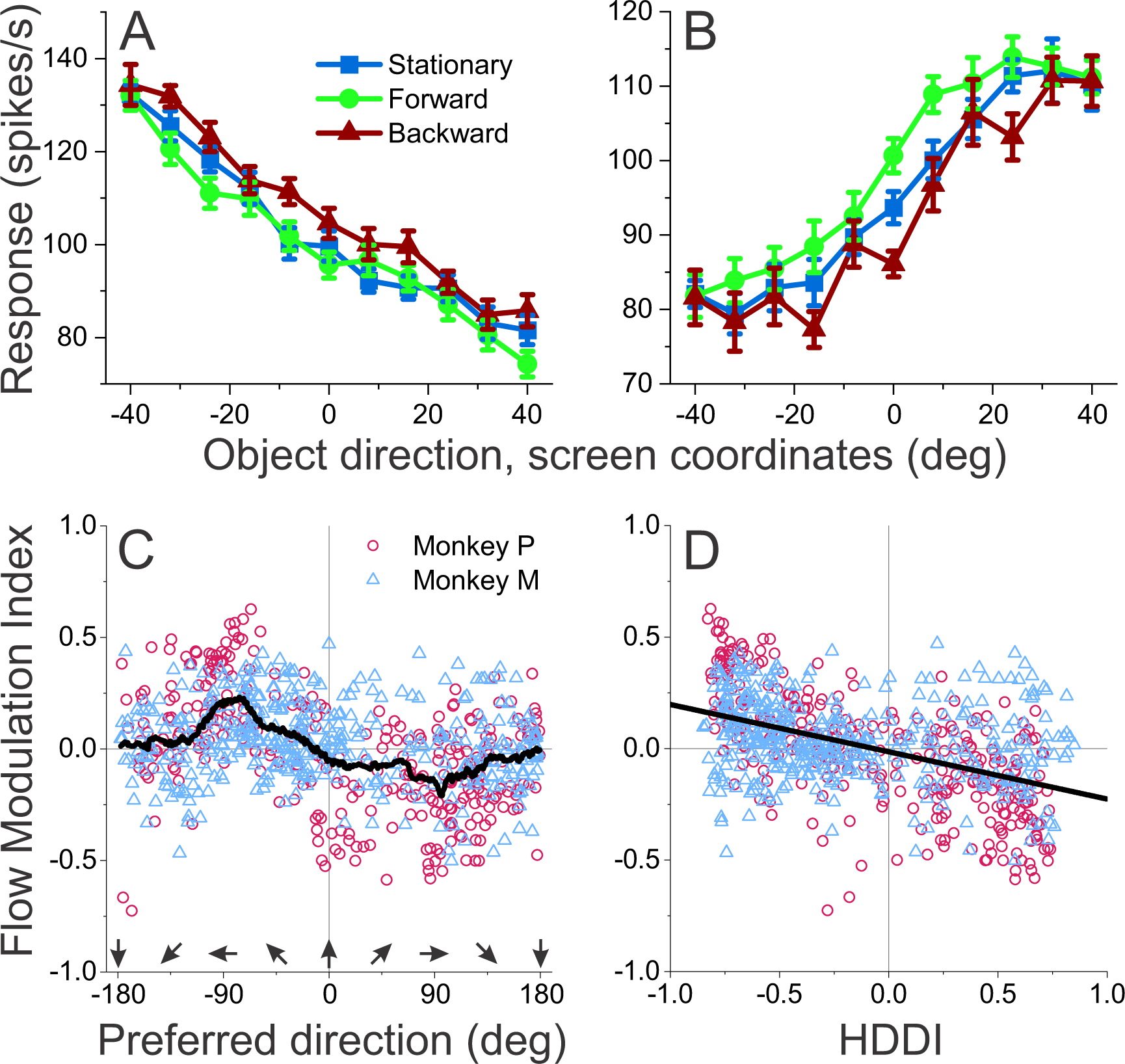
Modulation of MT firing rates by optic flow depends on direction tuning. **(A-B)** Firing rates of two units, recorded during the same session in which the object was in the left visual hemi-field. Data are shown separately for the stationary (blue squares), forward (green circles), and backward (red triangles) self-motion conditions. Error bars denote SEM. **(A)** For a unit that prefers leftward object motion, responses during backward self-motion are greater than responses during forward self-motion. **(B)** For a simultaneously recorded unit that prefers rightward object motion, responses during forward self-motion are greater than those during backward self-motion. **(C-D)** Flow-modulation index (FMI) across the population of 737 units depends systematically on aspects of direction tuning. Color and shape of symbols denote monkey identity: monkey M (teal triangles) and monkey P (purple circles). **(C)** FMI is circularly correlated with preferred direction, where a preference of 0 denotes the upward task direction reference. The black trace denotes a running median FMI, computed within a preferred direction window of 90° (data pooled across monkeys). **(D)** FMI is inversely correlated with selectivity for horizontal motion, as measured by the horizontal direction discrimination index (HDDI). Black line indicates the line of best fit (linear regression, data pooled across monkeys).

##### Dissociating effects of optic flow and choice on neural responses

FMI measures differences in response associated with self-motion direction indicated by optic flow. However, the perceptual biases induced by optic flow mean that choices are correlated with self-motion direction. Thus, it is possible that neural effects captured by FMI simply reflect a correlation of neural responses with choices. To dissociate the contributions of choice and self-motion direction to MT responses, we took advantage of the subset of object directions for which monkeys made both choices for each self-motion direction. This allowed a conditioning analysis to measure distinct effects of choice and self-motion direction, analogous to that used previously (Nogueira et al., 2017; Sasaki et al., 2020).

To quantify choice-related activity in individual units, we computed the well-established choice probability (CP) metric (Britten *et al*., 1996). For each distinct combination of object direction and self-motion direction (forward/backward), the distribution of responses was z-scored and then divided into two groups based on whether the animal made a leftward or rightward saccade to indicate their choice. Because this was done separately for each self-motion direction, effects of optic flow on the CP metric were removed. Z-scored responses were then pooled across unique stimulus conditions as long as there were at least 3 choices made toward each target location. ROC analysis was then applied to the pooled z-scores for the two choice groups, and CP was defined as the area under the ROC curve. For our purposes, CP was not referenced to each neuron’s preferred direction; rather CP > 0.5 corresponds to a preference for rightward choices and CP < 0.5 corresponds to a preference for leftward choices. This avoids potential issues with defining the “preferred” stimulus when choice effects are large (Zaidel et al., 2017).

We used an analogous ROC-based metric to quantify response modulations related to the direction of self-motion simulated by optic flow. This ‘flow probability’ (FP) metric is computed like CP, but swapping the roles of variables that represent choice (left vs. right) and self-motion direction (forward vs. backward). For each distinct combination of object direction and choice, responses were z-scored and sorted into two groups based on self-motion direction. If there were at least 3 trials for forward and backward self-motion, normalized responses from that condition were pooled with other conditions that met the same criteria. ROC analysis was applied to the pooled z-scores that were sorted by self-motion direction. The resulting metric was then multiplied by sign(*ObjXLocation*), such that it’s sign would be expected to match the sign of FMI if FMI is driven solely by effects of self-motion direction. Thus, FP provides a metric similar to FMI but removes any response modulations that depend on choice.

##### Time course analyses

To observe the temporal dynamics of MT responses to combined object motion and optic flow, we analyzed the time course of the normalized population response, FMI, and HDDI. Only units recorded during sessions with 2-s trials were included in the time course analyses (669/727 units). Data were pooled across monkeys.

First, we computed a peristimulus time histogram (PSTH) of the population response to vertical object motion. Separate PSTHs were calculated for each self-motion condition. Firing rates were calculated by taking a moving sum of the spike trains for a given self-motion direction with a window width of 50 ms. Each unit’s firing rates were normalized (across all self-motion conditions) so that the unit’s peak response was assigned a value of one. A PSTH was computed for each unit from these normalized responses, and the population PSTH was computed as the mean PSTH across units. Spontaneous firing rates were not subtracted when normalizing each unit’s responses, and the mean normalized baseline response was 0.28. To test for a difference in normalized response between stationary and self-motion trials, we performed a rank-sum test on the distributions of stationary and self-motion (combining forward and backward) responses for each of the 2501 time points from the onset of the stimulus to 500 ms after stimulus offset. A time point was considered to have a significant difference in response between stationary and self-motion conditions if the rank-sum test revealed a significant difference with a Bonferroni correction for multiple comparisons (*p* < 0.05/2501 = 2.00 × 10^-5^).

A time course of HDDI was computed by calculating HDDI for each unit in 50-ms bins. Units were grouped according to the cosine (horizontal component) of their preferred direction in intervals of 0.25, and the mean HDDI time course was computed for each group. Periods of significant directional responses were defined as time windows in which the distribution of HDDIs differed between the group of units that preferred rightward motion (*cos(preferred direction)* > 0.75) and the group that preferred leftward motion (*cos(preferred direction)* < -0.75). We used rank-sum tests to compare the distributions of HDDI between groups within each of the 51 time windows from stimulus onset to 500 ms after stimulus offset. A time window was determined to have a directional response if the rank-sum test reached significance with a Bonferroni correction for multiple comparisons (*p* < 0.05/51 = 9.80 × 10^-4^).

Similarly, a time course of FMI was computed by calculating FMI for each unit in 50-ms bins. Units were grouped according to HDDI in intervals of 0.2, and the mean FMI time course was calculated within each group. Periods of significant optic flow modulation were defined as time windows in which FMI among units with the strongest positive HDDI (HDDI > 0.6) differed from FMI among units with the strongest negative HDDI (HDDI < -0.6). Rank-sum tests compared the distributions of FMI between groups in each of the 51 time windows from stimulus onset to 500 ms after stimulus offset, and a Bonferroni correction for multiple comparisons (*p* < 0.05/51 = 9.80 × 10^-4^) was implemented to assess statistical significance.

#### Population decoding

##### Decoding procedure

To investigate how well area MT carries information about choice and object motion, logistic regression models were trained to classify each of these variables from a linear combination of the responses of a population of simultaneously recorded MT units. These linear classifiers were implemented in Matlab using the function ‘fitclinear’ with a logistic regression learner and 10-fold cross-validation. Classifiers assigned a weight to the activity of each unit, indicating how strongly the unit contributed to the decoder’s predictions along with its sign. The weights correspond to the components (one per unit) of a vector of decoding weights. Decoding results were robust to variations in the type of decoder used (logistic regression, support vector machine, Fisher linear discriminant), and the type of cross-validation (5-fold, 10-fold, none). Thus, we only present results for the logistic regression decoder with 10-fold-cross-validation.

A separate linear classifier was trained to decode each variable of interest. First, we trained a linear decoder to predict the monkey’s choice on each trial (*choice* decoder), with rightward choices coded as +1 and leftward choices coded as -1.

Second, we trained a classifier to decode stimulus direction in screen coordinates (*stimulus_screen_* decoder). For this decoder, the stimulus was coded as the sign of the object’s direction relative to vertical on the display: +1 for rightward object motion, -1 for leftward object motion. Since straight upward motion does not have a sign, trials in which the object moved straight upward were not included in training this model.

However, firing rates from these trials were used to predict stimulus direction once the decoder was trained. Finally, we trained a linear classifier to decode the stimulus in world coordinates (*stimulus_world_* decoder). The object direction in world coordinates was computed as the vector difference of the object direction on the screen and the optic flow vector that would have appeared at the (*x, y, z*) location of the center of the object. Positive values of object direction (rightward object motion in the world) were coded as +1, while negative values (leftward object motion in the world) were coded as -1.

In addition to these three main decoders, a variation of the stimulus decoders was trained only on trials in which no self-motion was simulated (stationary background dots), such that there can be no distinction between world and screed coordinates in the training set. The decoder was then tested on both stationary trials and trials with self-motion. Since this *stimulus_NoSM_* decoder did not have access to neural responses when there was forward/backward optic flow, it served as a control. We also trained an analogous decoder to classify the monkey’s choices based only on responses in the no self-motion condition (*choice_NoSM_* decoder). If this *choice_NoSM_* decoder could predict choice biases seen in behavior during the self-motion conditions, it would suggest that choice-related activity alone is sufficient to account for the effects of background optic flow on MT responses.

##### Analysis of decoding results

Trained linear classifiers predicted stimulus or choice direction on held out trials, using 10-fold cross-validation. The reports of the decoders were analyzed in the same manner as the behavioral data, by tabulating the proportion of “rightward” choices of the decoder and plotting this proportion as a function of stimulus direction in either world or screen coordinates. This produced a set of “psychometric” curves for each decoder, and PSE shifts and FP gains were computed from these decoder psychometric functions in the same way that behavior was analyzed. The FP gains from each decoder were compared to those from the other decoders and to the monkey’s behavioral FP gains. Spearman’s rank correlations were computed between pairs of decoders to determine whether the PSE shifts produced by two decoders covary across recording sessions. Wilcoxon signed-rank tests were used to assess whether two decoders produced PSE shifts (or FP gains) across sessions that come from distributions with different medians.

Multiple linear regression was used to determine what features of a unit’s tuning contribute to its individual decoding weight for each different type of decoder. The regressors were HDDI, to represent a unit’s strength of direction tuning, and FMI, to represent the strength of a unit’s modulation in the presence of optic flow. We took the components of the vector of decoding weights for a particular decoder, with each component corresponding to a single unit, and regressed them against the HDDI and FMI values for each unit. Prior to regressions, decoder weights were normalized within each session by computing their z-scores. Linear regressions were computed using Matlab’s fitlm function, after combining data across recording sessions for each animal. This yielded, for each type of decoder, a regression coefficient that captured the relationship between decoding weights and HDDI/FMI values, as well as a regression coefficient that reflected the interaction between decoding weights and HDDI/FMI values. Regression coefficients that are significantly different from 0 indicate a variable or interaction that plays a significant role in predicting the neural weights for that decoder.

## Results

Two macaque monkeys performed a fine discrimination of object motion direction in the presence of optic flow simulating forward or backward self-motion (Figure 1A,B; 20 sessions for monkey M, 19 sessions for monkey P). We first examine the perceptual biases that were induced by optic flow. We next describe the effect of optic flow on the responses of 727 units that were recorded from area MT during the discrimination task. Finally, we used within-session population decoding approaches to examine whether MT activity can account for the behavioral effects, and whether it can represent object motion in screen or world coordinates. Note that, since the fixation target remains centered on the screen during simulated self-motion, screen coordinates and retinal coordinates would be isomorphic if the eyes remain perfectly fixated during stimulus presentation. Since fixation cannot be perfect, we adopt the terminology of screen coordinates.

### Perceived object motion direction is biased in the presence of optic flow

The proportion of rightward choices, relative to vertical, was plotted as a function of object motion direction (in screen coordinates) to construct a psychometric function for each optic flow condition. Choices were biased in the presence of optic flow, and the direction of the bias depended on the location of the object (Figure 2 A,B). When the object was in the right visual hemi-field (Figure 2A), choices were biased leftward during simulated forward self-motion and rightward during backward self-motion. This effect is consistent with the flow-parsing hypothesis, as forward self-motion produces optic flow vectors in the right visual hemi-field with a rightward component, such that subtraction of these vectors is expected to produce a leftward perceptual bias (Fig. 1A, right panel). Backward self-motion, conversely, produces optic flow vectors in the right hemi-field with a leftward component, such that subtraction should induce a rightward bias (Fig. 1B, right panel). When the object to be discriminated was in the left visual hemi-field (Figure 2B), biases induced by optic flow were reversed, as expected from the flow-parsing hypothesis.

The effect of optic flow on perceived direction of object motion was quantified for each session by computing a PSE shift, which is the difference in PSE between forward and backward self-motion, multiplied by the sign of the object’s horizontal location in the visual field (Eqn. 3). When computed this way, a positive PSE shift always indicates a perceptual shift in the direction that is predicted by flow-parsing. The PSE shifts for the sessions in Figure 2A, B are 12.7° and 26.8°, respectively. For perfect flow parsing, the PSE shifts for these two sessions are expected to be 30° and 20°, respectively (interval between dashed vertical lines, Fig. 2 A, B).

Since stimulus conditions and expected PSE shifts varied across sessions and animals, we computed a flow-parsing gain (FP gain) as the ratio between the observed PSE shift and the expected PSE shift. The FP gain should be 1 if the monkey’s perceptual reports reflect complete subtraction of background optic flow, 0 if no flow-parsing occurs at all, and intermediate if the monkey’s behavior reflects partial subtraction of optic flow. FP gains may also be greater than 1 if the monkey overcompensates for self-motion.

The distribution of FP gains measured during 39 recording sessions (20 for monkey M, 19 for monkey P) is shown in Figure 2C. Median FP gains were 1.14 for monkey M and 0.32 for monkey P, and this difference was significant (Wilcoxon rank-sum test, *Z* = 4.5097, *p* = 6.493 × 10^-6^). While both monkeys’ FP gains were significantly greater than 0 (Wilcoxon signed-rank test, monkey M: *Z* = 3.920, *p* = 8.858 × 10^-5^; monkey P: *Z* = 3.823, *p* = 1.318 × 10^-4^), monkey P’s FP gains were significantly less than 1 (Wilcoxon signed-rank test, *Z* = -3.662, *p* = 2.502 × 10^-4^). This shows that monkey P exhibited partial flow-parsing, consistent with previous results in humans (Dokka et al., 2015; Fajen et al., 2013; Layton and Niehorster, 2019; Niehorster and Li, 2017). In contrast, monkey M’s median FP gain was marginally greater than 1 (Z = 1.904, p = 0.0569), suggesting some overcompensation for self-motion when judging object motion.

As documented previously (Peltier *et al*., 2020), FP gains for both monkeys decreased over time during training, likely due to the reward regimen used (see Discussion). For monkey M (Suppl. Fig. 1A), flow-parsing gains started well above 1, signifying a large overcompensation for self-motion, but decreased toward 1 by the time recording sessions commenced. For Monkey P (Suppl. Fig. 1B), FP gains started closer to 1 and decreased over time, stabilizing around ∼0.3. Interestingly, when the object was moved to the opposite hemi-field (Suppl. Fig. 1B, leftward-facing triangles), flow-parsing gains increased toward their original values and subsequently declined again. For both animals, FP gain were relatively stable during the period of time when recording sessions took place (colors in Suppl. Fig. 1).

### Effect of optic flow on MT responses depend on motion direction preference

We next examined whether a neural correlate of these perceptual biases is manifest in the activity of neurons in area MT. While monkeys performed the discrimination task described above, 24-channel linear electrode arrays were used to record activity from area MT (see Methods). In order to be included in analyses, units had to be directionally tuned, needed to have a receptive field that could be well-fit with a 2-dimensional Gaussian function, and were required to have less than 25% overlap between the receptive field and the background optic flow (see Methods for details). Of the 974 units recorded over 39 sessions, 727 units met these inclusion criteria (317 units from 19 sessions in monkey P, 410 units from 20 sessions in monkey M). The scarcity of well-isolated single units (18/727) did not allow us to analyze them separately from multi-units, so all units were pooled in the analysis. Average receptive fields for each recording session are shown in Suppl. Fig. 3.

Direction tuning curves of two example units, recorded during the same session of the discrimination task, are shown in Figure 3 A,B. In this session, the target object was in the left visual field. The tuning curve of a unit that preferred leftward object motion (Figure 3A) was modulated by optic flow such that backward optic flow enhanced firing rates relative to forward optic flow. The tuning curve of a unit that preferred rightward object motion (Figure 3B) shows the opposite pattern; firing rates were greater during forward self-motion than backward self-motion. Although these units were presented with the same optic flow patterns, they exhibited opposite effects of optic flow on responses to object motion. Crucially, the effects shown by both units are in the directions expected if optic flow shifts the population activity profile, as illustrated in Figure 1C.

We quantified response modulations caused by background optic flow by computing a flow modulation index (FMI, Eqn. 12). FMI is a normalized measure of the difference in response between forward and backward self-motion conditions, averaged across object motion directions. The sign of FMI is adjusted according to the location of the object relative to the fixation target (Eqn. 12), such that data could be combined across recordings in the left and right hemi-fields.

If flow parsing causes a shift of population activity in MT that explains perceptual biases (Figure 1C), the firing rates of individual units should be affected differently depending on their preferred directions. Specifically, FMI should be positive for leftward-preferring neurons and negative for rightward-preferring neurons; neurons preferring near-vertical directions should have FMI values closer to zero, as a shift of the population activity has less effect on these neurons (Figure 1C). Indeed, this pattern was observed across the populations of MT neurons recorded in each animal. Figure 3C demonstrates a roughly sinusoidal relationship between FMI and preferred direction, and a circular-linear correlation reveals a highly significant relationship for each animal (monkey M: *r_circular_* = 0.263, *p* = 6.96 × 10^-7^; monkey P: *r_circular_* = 0.611, *p* < 1 × 10^-14^). While there is clearly considerable variability in this relationship across the population of MT units, this pattern is consistent with a shift in the MT population response that could account for the perceptual biases induced by flow parsing.

As a complementary analysis, we would also expect FMI to depend on the strength of MT units’ selectivity for leftward vs. rightward motion, given that the task involves discriminating the horizontal component of motion. We quantified selectivity for leftward vs. rightward motion with a quantity called the horizontal direction discrimination index (HDDI, see Methods, Eqn. 11). A positive HDDI indicates a rightward direction preference, while a negative HDDI indicates a leftward preference. The magnitude of the HDDI indicates the strength of directional selectivity over the range measured during the discrimination task. FMI is plotted as a function of HDDI in Figure 3D, revealing the expected linear relationship. FMI and HDDI are strongly correlated (Spearman’s rank correlation, monkey M: *r_s_* = -0.244, *p* = 6.40 × 10^-7^; monkey P: *r_s_* = -0.706, *p* = 1 × 10^-48^), indicating that neurons that better discriminate horizontal motion are more affected by optic flow.

### Distinguishing effects of background optic flow from choice-related responses

We have shown that background optic flow modulates MT responses in a manner that depends systematically on the direction preference of neurons relative to the vertical discrimination boundary. However, since optic flow also biases choices of the monkeys, one possibility is that neural response modulations simply reflect choice-related activity in MT (Britten *et al*., 1996; Nienborg et al., 2012; Purushothaman and Bradley, 2005; Uka and DeAngelis, 2004), rather than a mechanism of flow parsing per se. Thus, it is unclear whether FMI primarily reflects effects of background optic flow, effects of choice, or a mixture of the two. To address this issue, we applied a method (Nogueira *et al*., 2017; Sasaki *et al*., 2020) for dissociating effects of stimulus context (background optic flow direction) and choice on neural responses (see Methods for details). In brief, for a subset of object directions that produced choices in both directions for each optic flow condition, we used a z-scoring and conditioning approach to measure the effect of optic flow direction on neural responses while conditioning on choice (flow probability, FP), and to measure the effect of choice on neural responses while conditioning on optic flow direction (choice probability, CP). If we see effects of either optic flow or choice after conditioning on choice and optic flow, respectively, then these effects cannot be accounted for by the conditioned variable.

Figure 4 A,B shows that flow probability (FP) exhibits a clear, systematic dependence on preferred direction (circular-linear correlation, monkey M: *r_circular_* = 0.323, *p* = 5.45 × 10^-10^; monkey P: *r_circular_* = 0.587, *p* < 1 × 10^-14^) and HDDI (Spearman’s rank correlation, monkey M: *r_s_* = -0.248, *p* = 3.83 × 10^-7^; monkey P: *r_s_* = -0.691, *p* = 4.4 × 10-^46^) that is very similar to the dependencies of FMI in Fig. 3C,D. In contrast, choice probability (CP) shows an inconsistent dependence on direction preference (circular-linear correlation, monkey M: *r_circular_* = 0.312, *p* = 1.56 × 10^-4^; monkey P: *r_circular_* = 0.108, *p* = 0.155) and exhibits a weak correlation with HDDI that has the opposite sign compared to FP (monkey M: *r_s_* = 0.119, *p* = 0.015; monkey P: *r_s_* = 0.125, *p* = 0.026). Further analysis shows that FP is strongly correlated with FMI, whereas CP is uncorrelated with FMI and weakly negatively correlated with FP (Suppl. Fig. 4). These findings demonstrate that background optic flow induces contextual modulations of MT responses that cannot be simply explained by choice-related activity.

**Figure 4.**
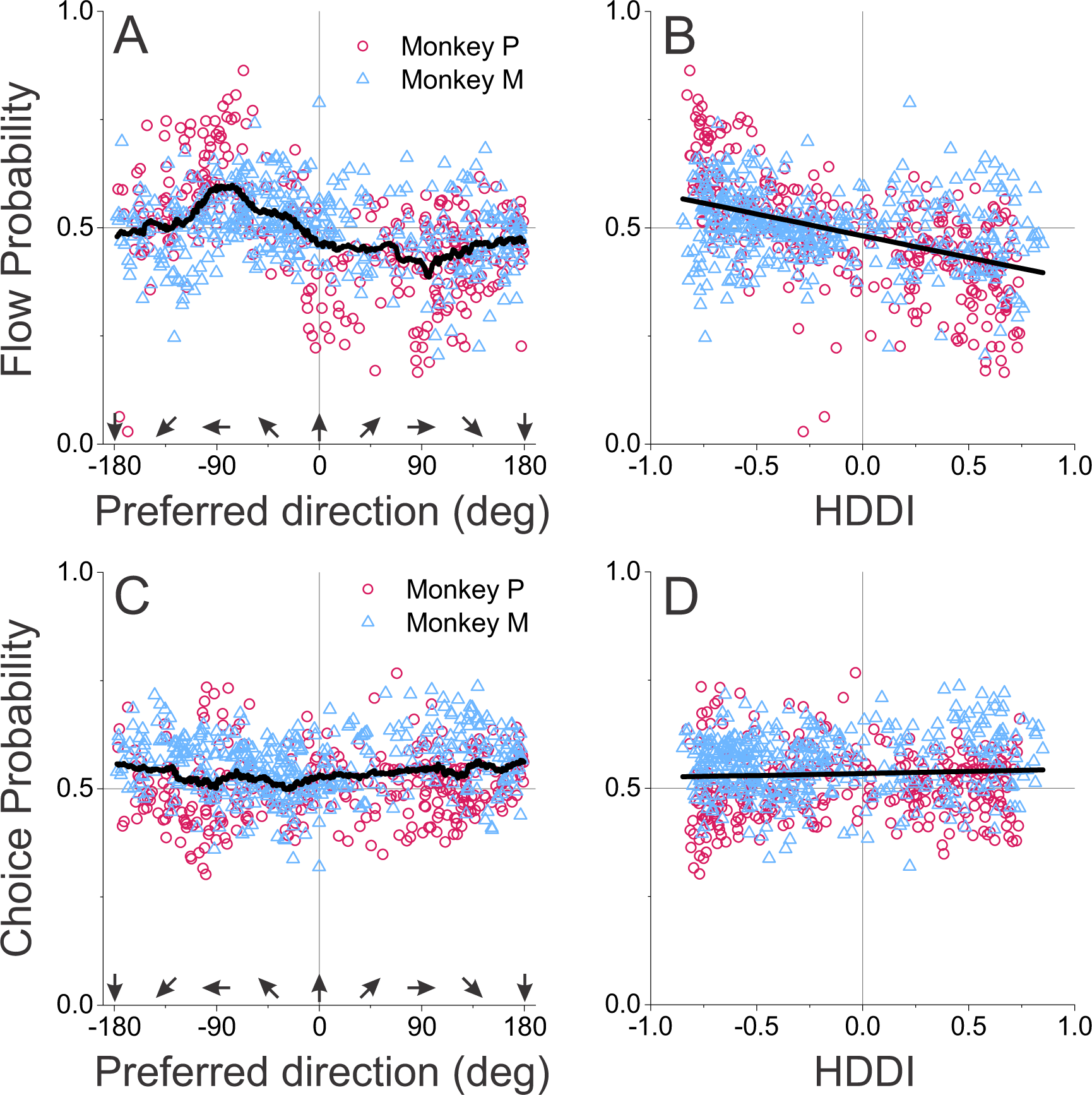
Effects of optic flow on MT responses are distinct from choice-related activity. Neural responses were analyzed to dissociate effects of background optic flow (Flow Probability, FP) from choice-related response modulations (Choice Probability, CP), as detailed in Methods. (A, B) Flow probability is robustly correlated with both neuronal preferred direction and HDDI, similar to the results for FMI (format as in Figure 3C,D). (C, D) In contrast, choice probability is not systematically related to either flow probability or HDDI. This reveals that effects of object flow background on MT responses are dissociable from choice-related modulations.

### Strength of optic flow modulation cannot be explained by surround suppression

One might expect that effects of optic flow on MT responses are related to interactions between an MT neuron’s classical receptive field and its inhibitory surround. For many MT neurons, responses to a stimulus in the classical receptive field are suppressed by motion in the surround (Allman et al., 1985), with suppression being strongest when the velocity of surround motion matches the cell’s preference. Could this form of directional surround suppression account for some or all of the observed effects of background optic flow on MT responses?

In our experimental design, we attempted to minimize contributions of surround suppression by masking out a region of the background flow field at least twice as large as the receptive field. Nevertheless, if masking was not completely effective, we might expect neurons with larger optic flow modulations to have stronger surround suppression. The absolute value of FMI is plotted as a function of surround suppression index (SSI) in Suppl. Fig. 5A. We performed a multiple linear regression of the absolute value of FMI onto SSI and monkey identity. We found no significant main effect of SSI (*t*(723) = -0.918, *p* = 0.359), as well as no interactive effect of SSI and monkey identity (*t*(723) = -1.35, *p* = 0.177), demonstrating that neurons with greater flow modulation do not generally have stronger surround suppression. The only significant effect was a main effect of monkey identity (β = -0.0623, *t*(723) = -4.42, *p* = 1.16 × 10^-5^), indicating that |FMI| values were significantly greater in monkey P than in monkey M. Treating each animal separately, there was no correlation between SSI and |FMI| for monkey P (Spearman rank correlation: *r_s_* = -0.0284, *p* = 0.615), and a significant negative correlation for monkey M (*r_s_* = -0.193, *p* = 8.31 × 10^-5^). This negative correlation indicates that units with stronger surround suppression had weaker flow modulation, which runs counter to the notion that flow modulation arises from surround suppression. Similar results were obtained when plotting FP against SSI (Suppl. Fig. 5B). The lack of positive correlation between SSI and FMI suggests that optic flow modulates MT responses via a mechanism that is distinct from conventional surround suppression, perhaps through feedback from higher-level areas that encode optic flow, such as MSTd or VIP (see Discussion).

### HDDI and FMI develop with different time courses

If neural correlates of flow parsing in area MT rely on feedback or some other mechanism that requires additional processing time, there may be a delay in the effect of optic flow on MT responses. To explore this issue, we analyzed the time courses of response properties of the MT population (Figure 5). Figure 5A shows the normalized population response to vertical object motion, computed using a moving 50-ms analysis window. Each unit’s firing rates were normalized (without subtracting spontaneous activity) such that the unit’s peak response (across all three self-motion conditions) was 1, and a population PSTH was then computed.

**Figure 5.**
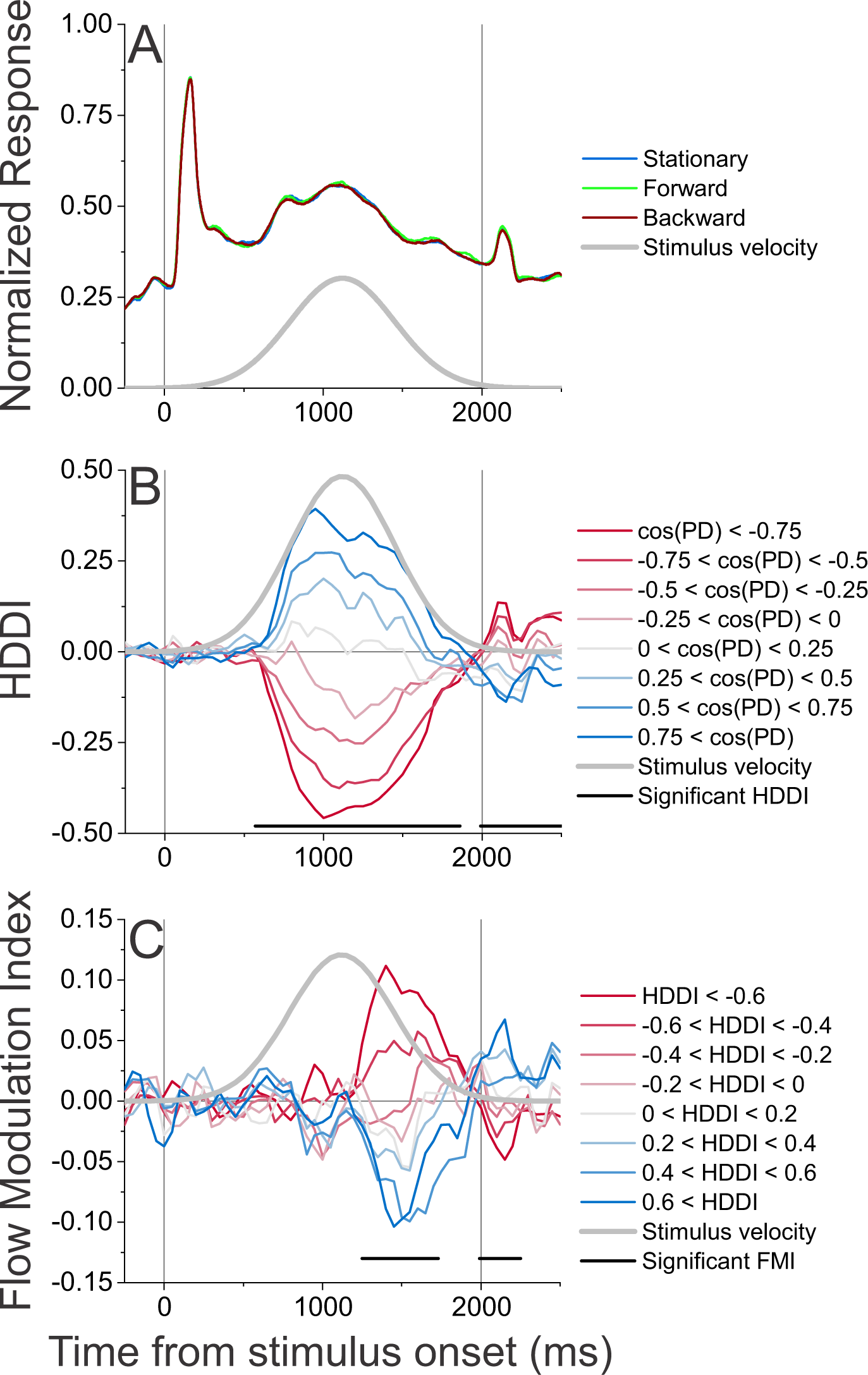
Time course of population response, HDDI, and FMI. In each panel, vertical lines indicate stimulus onset and offset, and the gray curve indicates the stimulus velocity profile. (A) Time course of normalized response to vertical (0 deg) object motion, averaged over all units. Color indicates self-motion direction; blue: stationary, green: forward, red: backward. (B) Mean HDDI time course for subsets of units, grouped according to the horizontal component of their preferred direction. Darker red curves indicate units with preferred directions closer to leftward (-90 deg), and darker blue curves indicate units with preferred directions closer to rightward (+90 deg). Horizontal black lines indicate time periods during which HDDI differs significantly between the darkest red and the darkest blue curves. (C) Mean FMI time course for subsets of units, grouped according to their HDDI values, where negative/positive HDDI values indicated a preference for leftward/rightward motion. Horizontal black lines indicate time periods during which FMI differs significantly between the darkest red and the darkest blue curves.

There was a robust transient response when dots appeared at the start of the trial, with a peak 160 ms after stimulus onset. This transient is presumably a response to the luminance onset of the dots, since motion begins sometime later. After the transient response, normalized response shows a broad central peak that roughly follows the temporal profile of stimulus velocity, peaking ∼1100 ms after stimulus onset. The time course of population response is practically indistinguishable between optic flow conditions (Fig. 5A, colored curves). Rank-sum tests revealed that there was no time point at which normalized responses differed significantly between the stationary and forward or backward self-motion conditions (*p* > 0.05; which is much greater than the threshold for significance after correction for multiple comparisons, which is *p* = 2.00 × 10^-5^). This indicates that optic flow did not elicit a net change in average response across the entire MT population. This likely occurs because forward and backward optic flow have opposite effects on MT units that prefer rightward and leftward directions (see Fig. 1C), and because rightward and leftward tuned neurons are roughly equal in number.

Figure 5B illustrates the time course of HDDI, with neurons grouped according to direction preference. As expected from the definition of HDDI, there is a clear separation in the HDDI time course between units that preferred rightward motion (shades of blue) and units that preferred leftward motion (shades of red). Approximately 600 ms after stimulus onset, units that preferred rightward motion developed positive HDDIs while units that prefer leftward motion developed negative HDDIs. This directionally selective response persists throughout most of the remainder of the stimulus period.

We identified periods of significant directional responses as time windows in which the distribution of HDDIs differed between the group of units that preferred rightward motion (*cos(preferred direction)* > 0.75; Figure 5B, darkest blue curve) and the group that preferred leftward motion (*cos(preferred direction)* < -0.75; Figure 5B, darkest red curve). We compared the distributions of HDDI between groups within each time window, with significance assessed by a Wilcoxon rank-sum test with Bonferroni correction for multiple comparisons, *p* < 9.80 × 10^-4^). There was a significant directional response from 600 to 1850 ms after stimulus onset, as well as a much weaker but significant reversal of selectivity in the ∼500 ms after stimulus offset. This reversal may be an adaptation effect, as shown previously following stimulus extinction (Kohn and Movshon, 2003; Van Wezel and Britten, 2002).

For comparison, Figure 5C illustrates the time course of FMI during the trial, with neurons grouped according to HDDI value. Strikingly, the time courses of FMI for rightward- and leftward-preferring units separate much later, during the second half of the visual stimulus presentation. We defined periods of significant optic flow modulation as time windows in which FMI for units with HDDI > 0.6 (Figure 5C, darkest blue curve) differed from FMI for units with HDDI < -0.6 (Figure 5C, darkest red curve). Rank-sum tests with Bonferroni correction for multiple comparisons (*p* < 9.80 × 10^-4^) revealed a main period of significant optic flow modulation ∼1300-1700 ms after stimulus onset. A weaker but significant reversal effect was also seen after stimulus offset, which may again be caused by response adaptation to the stimulus.

The fact that optic flow modulation of MT responses arose ∼700 ms after directional selectivity suggests that effects of optic flow on MT responses are mediated through additional pathways that are not necessary for the generation of direction selectivity. Flow modulation may be mediated through feedback from higher-level areas, such as MSTd or VIP, that respond selectively to optic flow (see Discussion).

### Single-session decoding of MT population responses

The fact that MT activity is modulated by background optic flow indicates that MT neurons represent more than just an object’s retinal velocity. Moreover, the dependence on direction preference (Fig. 3C,4A) is consistent with the hypothesis that background optic flow shifts the population hill of activity in the direction necessary to account for behavioral biases (Fig. 1C). However, it is unclear thus far whether the response modulations induced by optic flow are sufficiently strong to account for behavioral effects. To address this issue, we used logistic regression to perform within-session, trial-by-trial population decoding of MT responses. Decoders were trained to predict one of three variables: a *stimulus^screen^* decoder was trained to decode the direction of object motion in screen coordinates; a *stimulus_world_* decoder was trained to decode the direction of object motion in the world; and a *choice* decoder was trained to classify trials according to the monkey’s choice on each trial. In each case, decoders were trained, using 10-fold cross-validation, on data from all three self-motion conditions, and decoder performance was evaluated on held-out data (see Methods for details). Decoders were trained on spike counts computed over a time window from 1250-1750 ms after stimulus onset, as this time window revealed the largest FMI values (Figure 5C). The number of units included in decoding for each session ranged from 8 to 26, with a median of 19 units. For each type of decoder of MT responses, predicted psychometric curves, constructed from the choices generated by the decoder given the activity of the simultaneously recorded cells, were constructed for the 3 optic flow conditions, and FP gains were computed. This allows us to directly compare the FP gains of behavior with those predicted by the different decoders.

Psychometric functions from an example session for monkey P are shown in Figure 6A, and reveal a behavioral FP gain of 0.68. For this session, the stimulus_world_ decoder (Figure 6B) predicts a largely similar pattern of biases, with a predicted FP gain of 0.48. Thus, decoding of just a small population of MT units (N=21 in this case) can account for most of the behavioral biases and overall sensitivity of the monkey’s performance in this session. By comparison, the stimulus_screen_ decoder (Figure 6C) predicts much smaller PSE shifts, corresponding to a FP gain of 0.10. Thus, the same population of MT neurons can be decoded to obtain reasonable estimates of motion in either screen coordinates or world coordinates.

**Figure 6.**
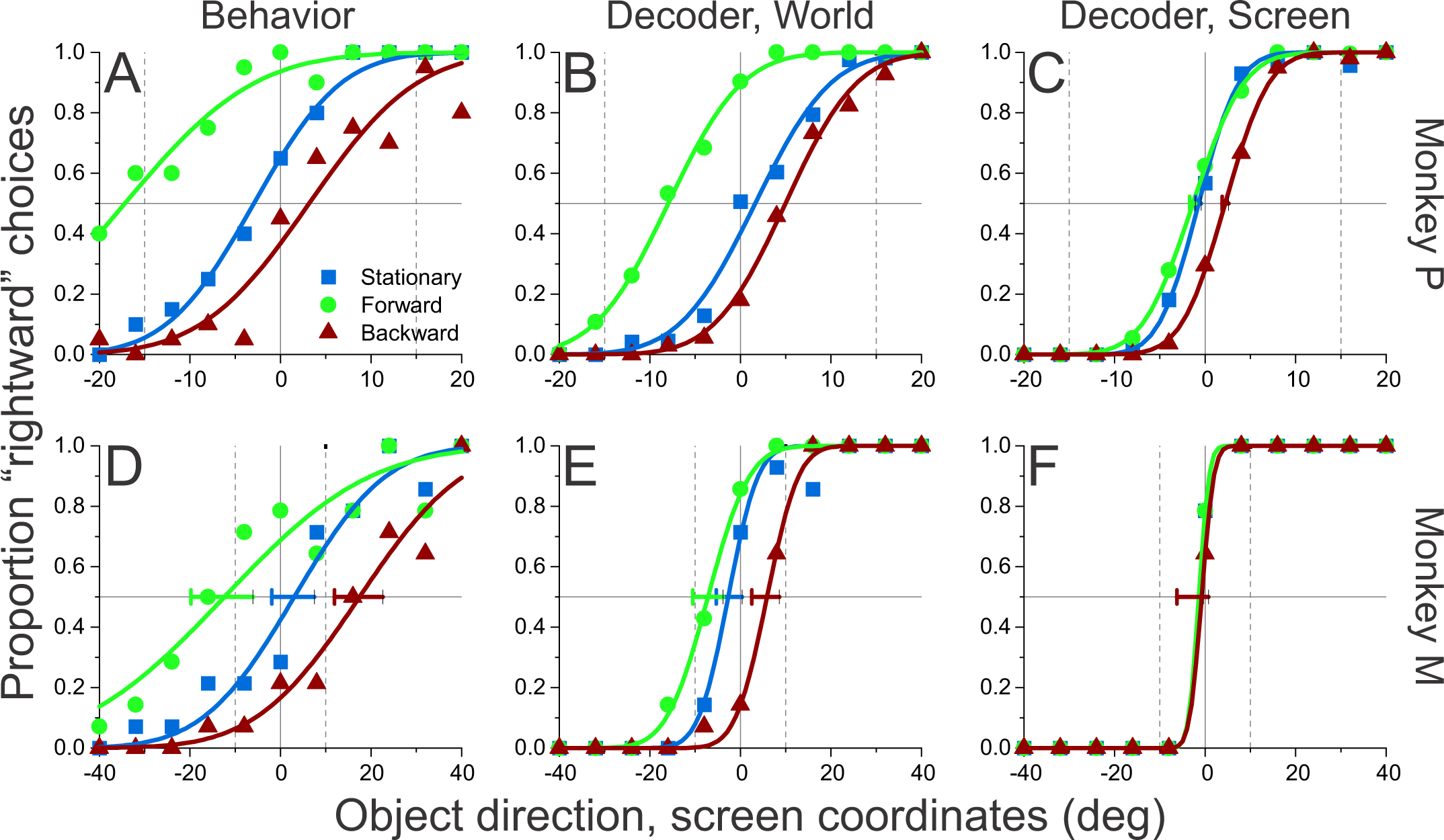
Psychometric functions representing monkey behavior and population decoder performance for two example sessions. Format as in Figure 2A,B. Vertical lines indicate the expected PSEs for complete flow-parsing. **(A)** Psychometric function reflecting one session of monkey P’s direction discrimination performance for an object located in the left visual hemifield. **(B)** Predicted psychometric function produced by the stimulus_world_ decoder, which was trained to discriminate object direction in world-centered coordinates from neural responses in the same for which the behavioral data are shown in panel A. **(C)** Psychometric function produced by the stimulus_screen_ decoder, which was trained to discriminate object direction in retinal coordinates (same session as panels A, B). **(D-F)** Psychometric data and decoder performance for one example session from monkey M.

Figure 6D-F shows similar results for an example recording session from monkey M (N=22 units). In this case, the FP gains are 1.48 for behavior, 0.65 for the stimulus_world_ decoder and 0.03 for the stimulus_screen_ decoder. The diversity of effects of optic flow on MT responses (Fig. 3C, 4A) likely allows units to be weighted differently between decoders to produce PSE shifts that vary according to the decoded variable, thus counterbalancing biases if there are systematic shifts in the tuning curves.

### Comparison between behavioral effects and decoder performance

For the stimulus_world_ decoder, Fig. 7A summarizes flow parsing effects across sessions for each animal. The median FP gains of the stimulus_world_ decoder are 0.45 for monkey M and 0.35 for monkey P, which are not significantly different from each other (Wilcoxon rank-sum test, *Z* = -1.03, *p* = 0.31). Pooling FP gains across monkeys, the median FP gain (0.41) is significantly less than 1 (Wilcoxon signed-rank test, *Z* = -5.43, *p* = 5.68 × 10^-8^) and significantly greater than zero (*Z* = 5.42, *p* = × 10^-8^), indicating that the stimulus_world_ decoder does not perfectly represent object motion in world coordinates for either animal.

**Figure 7.**
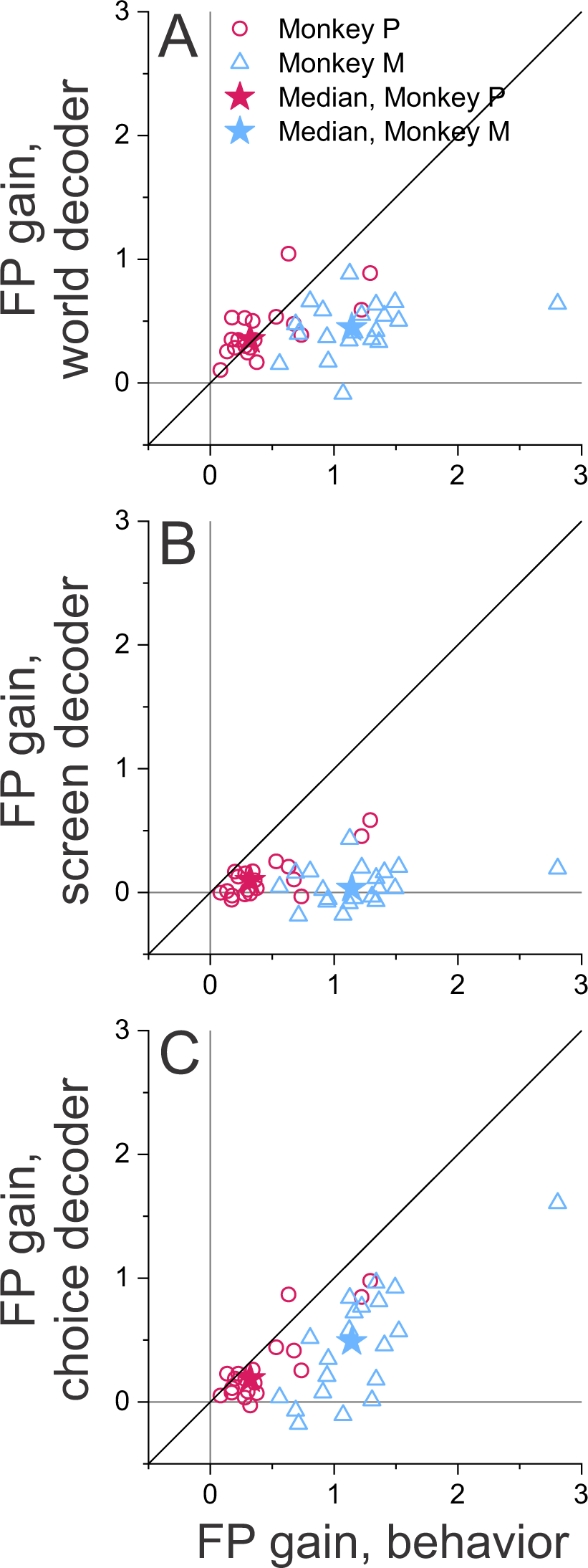
Summary of comparison between monkey behavior and population decoder performance. Each datum represents one experimental session from monkey M (teal triangles) or monkey P (purple circles). Star-shaped symbols indicate the median perceptual and decoder FP gains across sessions for each animal. (A) FP gains of the stimulus_world_ decoder are plotted against the monkeys’ perceptual FP gains. (**B**) FP gains of the stimulus_screen_ decoder are plotted against the monkeys’ perceptual FP gains. **(C**) FP gains of the choice decoder plotted against the monkeys’ perceptual FP gains.

Given that behavioral FP gains differed subtantially between monkeys, perhaps the more relevant question is whether performance of the stimulus_world_ decoder can account for behavioral performance, as summarized in Fig. 7A. For monkey P, there is no significant difference between median FP gains for behavior and the stimulus_world_ decoder (Wilcoxon signed-rank test, *Z* = -0.16, *p* = 0.87), indicating that neural effects of flow parsing in MT are sufficient to account for behavioral biases in this animal. Moreover, there is a significant correlation across sessions between FP gains for behavior and the stimulus_world_ decoder for monkey P (Spearman rank correlation, *r_s_* = 0.56, *p* = 0.014), indicating that variations in the neural effects account for some of the variations in behavior across sessions. By contrast, for monkey M, the median behavioral FP gain is significantly greater than the median FP gain of the stimulus_world_ decoder (Wilcoxon signed-rank test, *Z* = 3.92, *p* = 8.86 × 10^-5^), and there is no significant correlation between these measures across sessions (*r_s_* = 0.3098, *p* = 0.1835). Thus, while a linear decoder of stimulus_world_ predicts a partial shift (∼40%) toward a world-centered reference frame for both animals, this neural effect was able to fully account for behavioral biases in monkey P, but not in monkey M (see Discussion).

We also examined the performance of a decoder that attempts to classify object motion direction in screen coordinates. If this stimulus_screen_ decoder performs perfectly, it should yield FP gains equal to 0, regardless of whether perception is biased by optic flow. Median FP gains for the stimulus_screen_ decoder were 0.032 for monkey M and 0.098 for monkey P, and these values did not differ significantly between animals (Wilcoxon rank-sum test, Z = 1.05, p = 0.29). Pooled across animals, the median FP gain was 0.0324, which is significantly greater than 0 (Wilcoxon signed-rank test, *Z* = 2.71, *p* = 6.78 × 10^-3^). FP gains for the stimulus_screen_ decoder were significantly less than behavioral FP gains for both animals (Wilcoxon signed-rank test, monkey M: *Z* = 3.92, *p* = 8.85 × 10^-5^; monkey P: *Z* = 3.82, *p* = 1.32 × 10^-4^; Figure 7B). We found a modest but significant correlation between FP gains for behavior and the stimulus_screen_ decoder for monkey P (Spearman rank correlation: *r_s_* = 0.54, *p* = 0.02) but not for monkey M (*r_s_* = 0.33, *p* = 0.15).

FP gains significantly greater than zero for the stimulus_screen_ decoder suggest that effects of optic flow on MT responses are sufficiently pervasive that they cannot completely be discounted by a linear decoder to estimate motion in screen coordinates (although it is certainly possible that the stimulus_screen_ decoder would perform more ideally if based on larger populations of neurons than we recorded within each session). We observed similar effects when the decoder of motion direction was trained only on trials without background optic flow (stimulus_NoSM_ decoder). Because the median FP gains of the stimulus_NoSM_ decoder did not differ between animals (Wilcoxon rank-sum test, *Z* = -0.58, *p* = 0.56), we again pooled data across animals. The median pooled FP gain (0.10) of the stimulus_NoSM_ decoder was significantly greater than 0 (Wilcoxon signed-rank test, Z = 2.51, p = 0.012), indicating that this result was not simply driven by other variables (e.g., choice) that may covary with optic flow conditions.

Finally, we also trained a decoder to predict the choices of the animal, rather than stimulus direction. We found that the median FP gain for behavior was significantly greater than the median value for the choice decoder (monkey M: *Z* = 3.92, *p* = 8.86 × 10^-5^; monkey P: *Z* = 3.06, *p* = 0.002; pooled: *Z* = 5.18, *p* = 2.25×10^-7^). MT activity accounted for 58.3% of the behavioral effect for monkey P and 42.4% for monkey M. FP gains were also significantly correlated across sessions between behavior and the choice decoder (monkey M: *r_s_* = 0.65, *p* = 0.0025; monkey P: *r_s_* = 0.64, *p* = 0.0040), suggesting that MT activity can account for a substantial fraction of session-to-session variability in perceptual FP gains. Because choices are strongly related to the optic flow condition in behavior, the predictive capacity of the choice decoder could rely on either response modulations that are associated with choice or with background optic flow. To gain further insight, we trained a classifier to decode choice from just trials without self-motion (choice_NoSM_ decoder), as this condition affords the decoder with choice-related signals but not signals related to optic flow condition. This choice_NoSM_ decoder yields median FP gains (monkey M: -0.021; monkey P: 0.098) that are significantly less than those of the choice decoder trained on all conditions (Wilcoxon signed-rank test, monkey M: *Z* = 3.77, *p* = 1.63 × 10^-4^; monkey P: *Z* = 2.01, *p* = 0.044). These findings suggest that performance of the choice decoder is mainly driven by effects of background optic flow on MT responses, and this finding is consistent with the analysis of Figure 4, which shows that flow probability, but not choice probability, is systematically related to the direction preferences of MT neurons.

### Relationships between decoding weights and neural selectivity

The fact that these decoders can be trained to represent different variables suggests that different weights are assigned to the units depending on the decoded variable. To gain further insight into how neurons with different properties contribute to decoder performance, we used linear regression to examine relationships between decoding weights, HDDI, and FMI. The stimulus_screen_ decoder should place more weight on units with strong direction selectivity in screen coordinates (high magnitude of HDDI). However, because the computation of object motion in screen coordinates does not rely on information about optic flow, the stimulus_screen_ decoder should not selectively weight units according to FMI. A multiple regression analysis of the stimulus_screen_ decoder’s weights, w_StimulusScreen_, revealed that HDDI was significantly predictive of neuronal decoding weights (monkey M: β = 0.902, *t*(406) = 10.13, *p* = 1.188 × 10_-21_; monkey P: β = 0.726, *t*(311) = 6.33, *p* = 8.725 × 10^-10^), but FMI was not (monkey M: *t*(406) = -1.04, *p* = 0.299; monkey P: *t*(311) = 0.72, *p* = 0.472). This finding is consistent with the expectation that performance of the stimulus_screen_ decoder should not depend on modulations by background optic flow.

In contrast, computing object motion in world coordinates requires information about optic flow. Therefore, the stimulus_world_ decoder should significantly weight units based on the magnitude of both HDDI and FMI. Indeed our multiple regression analysis revealed that FMI was highly predictive of decoder weights, w_StimulusWorld_, for both animals (monkey M: β = -1.437, *t*(406) = -6.87, *p* = 2.38 × 10^-11^; monkey P: β = -1.871, *t*(311) = -8.26, *p* = 4.39 × 10^-15^). HDDI was significantly predictive of w_StimulusWorld_ for one animal (monkey M: β = 0.7571, *t*(406) = 8.732, *p* = 6.609 × 10^-17^) but not the other (monkey P: *t*(311) = -0.0909, *p* = 0.9277). There was a marginally significant interaction effect between HDDI and FMI on w_StimulusWorld_ for monkey M (β = 0.709, *t*(406) = 1.840, *p* = 0.066) but not for monkey P (*t*(311) = -0.616, *p* = 0.539). The significant contribution of FMI to decoding weights for the stimulus_world_ decoder demonstrates that units with strong response modulations by optic flow contribute to representing object motion in world coordinates. Very similar results were obtained using FP instead of FMI in this analysis, as expected given the strong correlation between FP and FMI measures (Suppl. Fig. 4A). In contrast, when FMI was replaced with CP in this analysis, there were no significant contributions (p>0.5) of CP to predicting either w_StimulusScreen_ or w_StimulusWorld_. This analysis further demonstrates that the capacity of MT population responses to predict perceptual biases associated with optic flow does not simply rely on choice-related activity.

## Discussion

Flow parsing has been proposed as a strategy for solving a crucial problem in vision: discounting the visual consequences of self-motion in order to compute object motion relative to the world (Rushton and Warren, 2005; Warren and Rushton, 2007; 2009a). Despite extensive psychophysical evidence for flow parsing in humans, the neural mechanisms underlying this process were unknown. We demonstrate here a neural basis for flow parsing in macaque area MT. Responses of MT units are modulated by surrounding optic flow in a systematic manner that depends on preferred direction. This effect is consistent with the hypothesis that optic flow shifts the population profile of activity in MT, leading to the observed behavioral biases. Crucially, these effects cannot be explained by choice-related activity or conventional surround suppression, as discussed further below. Consistent with the effects seen in individual units, single-session population decoding predicts biases in the same direction as seen in behavior, although weaker in magnitude. Together, these findings provide the first evidence of a mechanism for flow parsing at the level of single neurons and neural population activity.

### Systematic effects of optic flow on MT responses

We hypothesized that MT might reflect flow parsing via a shift of the population hill of activity as illustrated in Fig. 1C. Such a shift predicts that the response modulations induced by optic flow should depend systematically on direction preference relative to the reference direction of the task (which is vertical here). Indeed, our data (Fig. 3C, 4A) confirm this prediction, indicating that responses in area MT can account, at least partially, for the biases observed in behavior. However, results were variable, as many units showed little to no effect of optic flow. This variability suggests that a pure shift of the population hill of activity is likely too simple an explanation. Rather, some subpopulations of MT units may shift their representation of object motion toward world coordinates while others do not. Such diversity might enable MT to represent object motion in more than one coordinate frame, allowing for flexible readout according to task demands. In future work, it would be interesting to determine whether MT neurons with weak and strong modulations by optic flow have different projection targets.

For MT units that are modulated by optic flow, these changes typically manifest as differences in response between forward and backward optic flow that are fairly consistent across object motion directions (e.g., Fig. 3A,B). One possible explanation is that optic flow shifts direction-tuning curves in a manner consistent with representing object motion in the world. Another possibility is that MT responses are gain modulated by optic flow, such that optic flow acts to multiplicatively scale MT tuning curves. Due to the limited range of object directions presented in this experiment, our data do not allow us to clearly distinguish between these possibilities. In an ongoing study, we are measuring the full direction tuning curves of MT units in the presence of optic flow, such that we can better quantify how optic flow acts to modulate MT responses.

### Potential confounds of surround suppression and choice-related activity

Our findings demonstrate that optic flow surrounding a target object modulates neural responses in area MT and biases reports of motion direction. Here, we address two potentially less interesting explanations for these effects: non-classical surround suppression and choice-related modulations.

It is well established that receptive fields of many MT neurons have a non-classical surround that suppresses responses when surrounding stimuli have similar properties to the stimulus that activates the neuron (Allman *et al*., 1985; Born, 2000; Bradley and Andersen, 1998; DeAngelis and Uka, 2003). To limit involvement of the conventional surround, we employed a mask that was typically at least 2-fold larger than the classical receptive field. We also excluded units whose receptive fields overlapped with the surrounding optic flow (see Methods). Nevertheless, it is still possible that our findings could be a result of surround suppression. To assess this, we measured the size tuning of MT units, and we found no significant correlation between the magnitude of either FMI or FP and a quantitative measure of surround suppression (Suppl. Fig. 5). Thus, conventional surround suppression cannot account for our findings.

Another potential confound is choice-related activity. It is well-established that responses of MT neurons are weakly correlated with perceptual decisions (Britten *et al*., 1996; Purushothaman and Bradley, 2005; Uka and DeAngelis, 2004); as such, it is possible that the response modulations observed in MT simply reflect an effect of optic flow on choices, which subsequently modulates MT responses. To address this issue, we adapted an analysis (Sasaki *et al*., 2020) to isolate the effects of choice from the effects of background optic flow in the response of MT neurons (see Methods, Fig. 4, and Suppl. Fig. 4). This demonstrates clearly that flow-related modulations (Fig. 4 A, B), but not choice-related modulations (Fig. 4 C, D), are systematically related to direction preferences in MT. In addition, we demonstrate that decoding weights for the stimulus_world_ decoder are correlated with both the FMI and FP values of individual units, whereas these decoding weights are not correlated with CP values. Thus, our findings cannot be simply explained by choice-related activity, and instead suggest a modulation of MT responses that is specific to the inferred optic flow vector at the location of the target object.

### Comparison of behavioral and decoder biases

During recording sessions, the two animals had different magnitudes of behavioral effects. While monkey M showed a flow-parsing gain close to unity, monkey P had biases that were substantially smaller than predicted by perfect flow parsing (Fig. 2C). The incomplete compensation for self-motion exhibited by monkey P has been observed previously in humans (Dokka *et al*., 2015; Fajen *et al*., 2013; Layton and Niehorster, 2019; Niehorster and Li, 2017). In addition, both monkeys demonstrated a substantial decrease in flow-parsing gains over time, mainly during the training period prior to commencement of recordings (Suppl. Fig. 1, see also (Peltier *et al*., 2020)). For monkey P, flow-parsing gain decreased from near unity toward values < 0.5. In contrast, monkey M had flow-parsing gains that were initially much greater than unity, declined sharply toward unity prior to recordings, and declined gradually during recordings such that the final values were less than unity. A likely explanation for this reduction in flow-parsing gains over time is the variable reward scheme that we used (see (Peltier *et al*., 2020) for further discussion). Ideally, one would reward animals around their intrinsic perceptual biases, such that there is not pressure to decrease perceptual biases to maximize reward. In ongoing work, we have developed a Bayesian adaptive technique to estimate perceptual biases online, but this was not available when the present study was conducted.

For decoders trained to classify object direction in world coordinates, we found that single-session decoding in both animals predicted flow-parsing gains near 0.5, on average (Fig. 7A). For monkey P, decoder performance was sufficient to account for the flow-parsing gains seen in behavior; in contrast, for monkey M, decoder effects were substantially weaker than behavioral effects (Fig. 7A). It is possible that flow-parsing acts at multiple stages of motion processing, as suggested by human neuroimaging studies (Field et al., 2020; Kozhemiako et al., 2020; Pitzalis et al., 2020), and that effects on areas downstream of MT account for the larger behavioral effects in monkey M. However, it is also possible that decoding from larger neural populations in area MT could predict the larger biases seen in monkey M.

It may seem surprising that the visual system would compensate for self-motion at a relatively early processing stage such as area MT, which has often been thought to represent retinal image motion. However, it is important to point out that retinal motion information is not lost in MT. Due to the diversity of effects of background optic flow in MT units, population responses could be weighted differently to read out motion in either world or screen coordinates. Indeed, we find that decoding weights are correlated with FMI values for the stimulus_world_ decoder but not the stimulus_screen_ decoder. Correspondingly, we find that decoders trained to recover direction in screen coordinates are much less biased by optic flow (Fig. 7B). Thus, the representation we observe in MT may allow downstream areas to flexibly decode object motion in different reference frames.

Furthermore, flow parsing should only be done if an observer infers that there is an independently moving object in the scene. This was generally the case in our study because recording sites were selected to have receptive fields located closer to the horizontal meridian. Since the direction reference for the task was always vertical, optic flow vectors at the location of the target patch were typically close to orthogonal to the task reference. Recent psychophysical work shows that motion perception can transition from integration to segmentation depending on the similarity of an object’s motion to the background (Shivkumar et al., 2023). Thus, it is possible that flow parsing effects in MT would be modulated by causal inference regarding the likelihood of independent object motion, which is an ongoing topic of exploration in the laboratory.

### Potential neural networks of flow-parsing

As discussed above, decoding results from MT typically account for less than the full behavioral effects of flow parsing, suggesting that additional contributions to flow parsing may arise downstream of MT. Relevant to this, flow modulation effects were substantially slower (∼700 ms) to develop than directional responses (Fig. 5 B, C).While the reasons for such a long delay are unclear, it may reflect computations that are done downstream and then fed back to MT to modulate population responses. MT is known to receive feedback from areas that are involved in computing heading, such as areas MSTd (Bradley et al., 1996; Duffy and Wurtz, 1995; Fetsch et al., 2012; Gu et al., 2008; Gu *et al*., 2006) and VIP (Bremmer et al., 2002; Chen et al., 2011; 2013; Zhang and Britten, 2004; 2010). Another downstream area of interest is area V6, which is reported to contain subpopulations of neurons that represent motion in head-centered reference frames (Fattori et al., 2009; Galletti et al., 1995). The origin of the long delay of neural flow parsing effects in MT deserves further study. It is worth noting that other previous studies have examined how rotational optic flow modulates MT responses, and those effects were not found to have such a long delay (Kim et al., 2015; 2017).

Computational models of heading perception (Hatsopoulos and Warren, 1991; Layton and Browning, 2014; Layton and Fajen, 2016a; Layton et al., 2012; Perrone, 1992; 2012; Perrone and Stone, 1994; 1998; Royden, 1997) and flow parsing (Layton and Fajen, 2016b; Layton and Niehorster, 2019; Royden and Holloway, 2014) have primarily focused on the relationship between MT and MSTd. These models consist of a layer of small MT-like operators that are selective for motion direction and speed. These operators transmit their responses to a layer of large MSTd-like templates, whose receptive fields are built from the combination of MT operators with receptive fields spanning the template. The estimated heading is typically the preferred heading of the most active MSTd template, or a combination of the most active templates. In a model that was specifically designed to implement flow parsing (Layton and Fajen, 2016b), feedback from the MSTd layer to the MT layer enhances the activity of MT units with a preferred direction that disagrees with the most active MSTd template. It is currently unclear whether this type of model is consistent with our physiological findings or not. However, simultaneous recordings from MT and MSTd may be valuable for testing specific predictions of this type of model. Furthermore, it would be valuable to examine how reversible inactivation of MSTd affects flow-parsing behavior as well as the response modulations of MT neurons. Having established a starting point for understanding the neural computations of flow parsing, subsequent studies can be targeted to unraveling the relevant neural circuits.

## Supporting information

Stimulus video

## Acknowledgments

We thank Johnny Wen and Amanda Yung for programming assistance, as well as Dina Graf, Emily Murphy, Yinghui Qiu, and Victor Kogan for assistance with animal care and training. This work was supported by NEI R01 grant EY016178, NINDS U19 grant NS118246, and by the Computing Module of an NEI Core grant (EY001319).

---

**Supplementary Figure 1.**
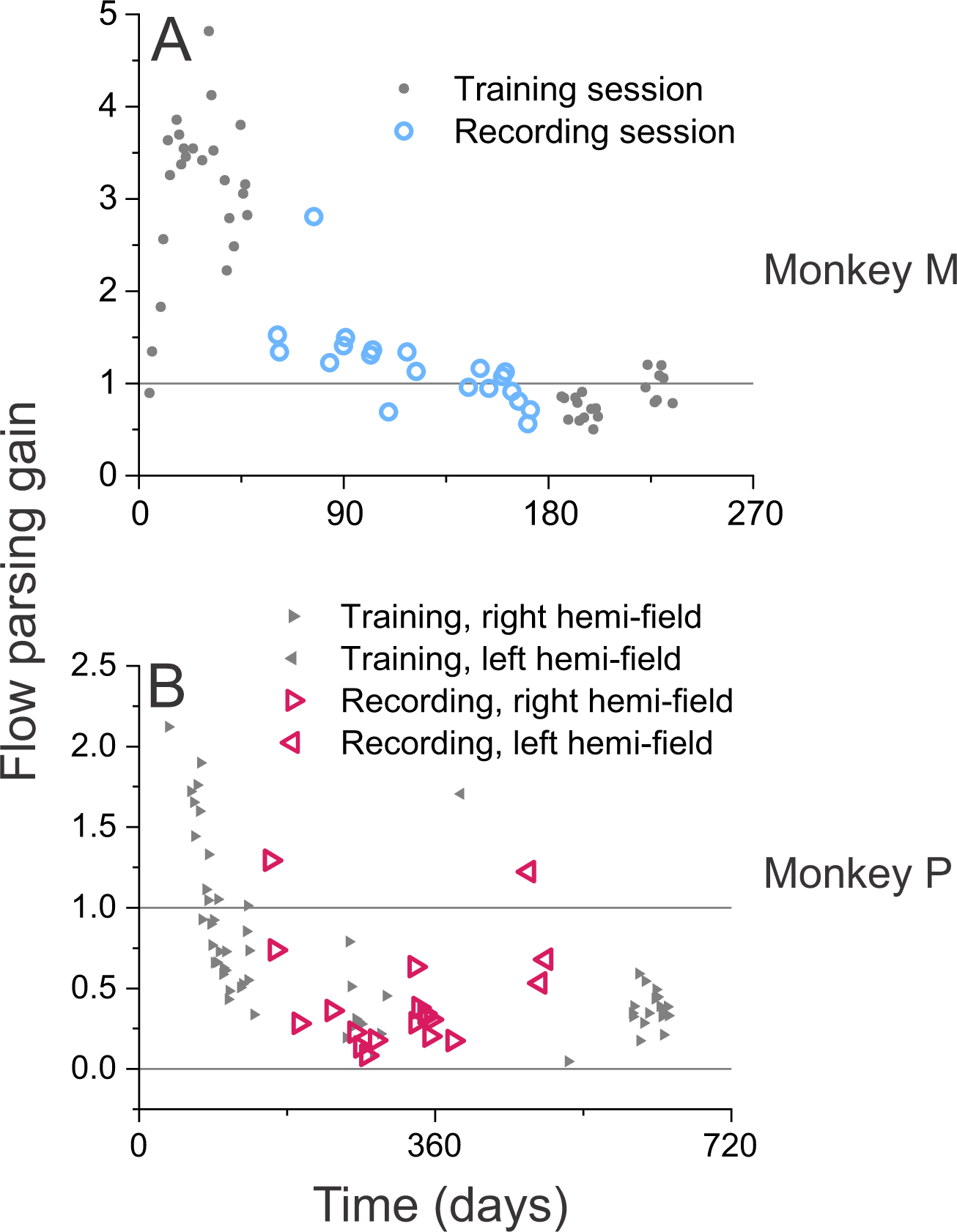
Flow-parsing gains gradually decreased over time. **(A)** Monkey M’s flow-parsing gains throughout 20 recording sessions (large, teal open circles), in addition to the flow-parsing gains over 44 training sessions (small, gray filled circles). Each gray circle indicates the mean flow-parsing gain for a training session that took place before or after recordings. **(B)** Monkey P’s flow-parsing gains throughout 19 recording sessions (large, purple open triangles), as well as flow-parsing gains over 62 training sessions (small, gray filled triangles). Right-facing triangles denote sessions in which the object was in the right hemi-field, and left-facing triangles denote sessions in which the object was in the left hemi-field.

**Supplementary Figure 2.**
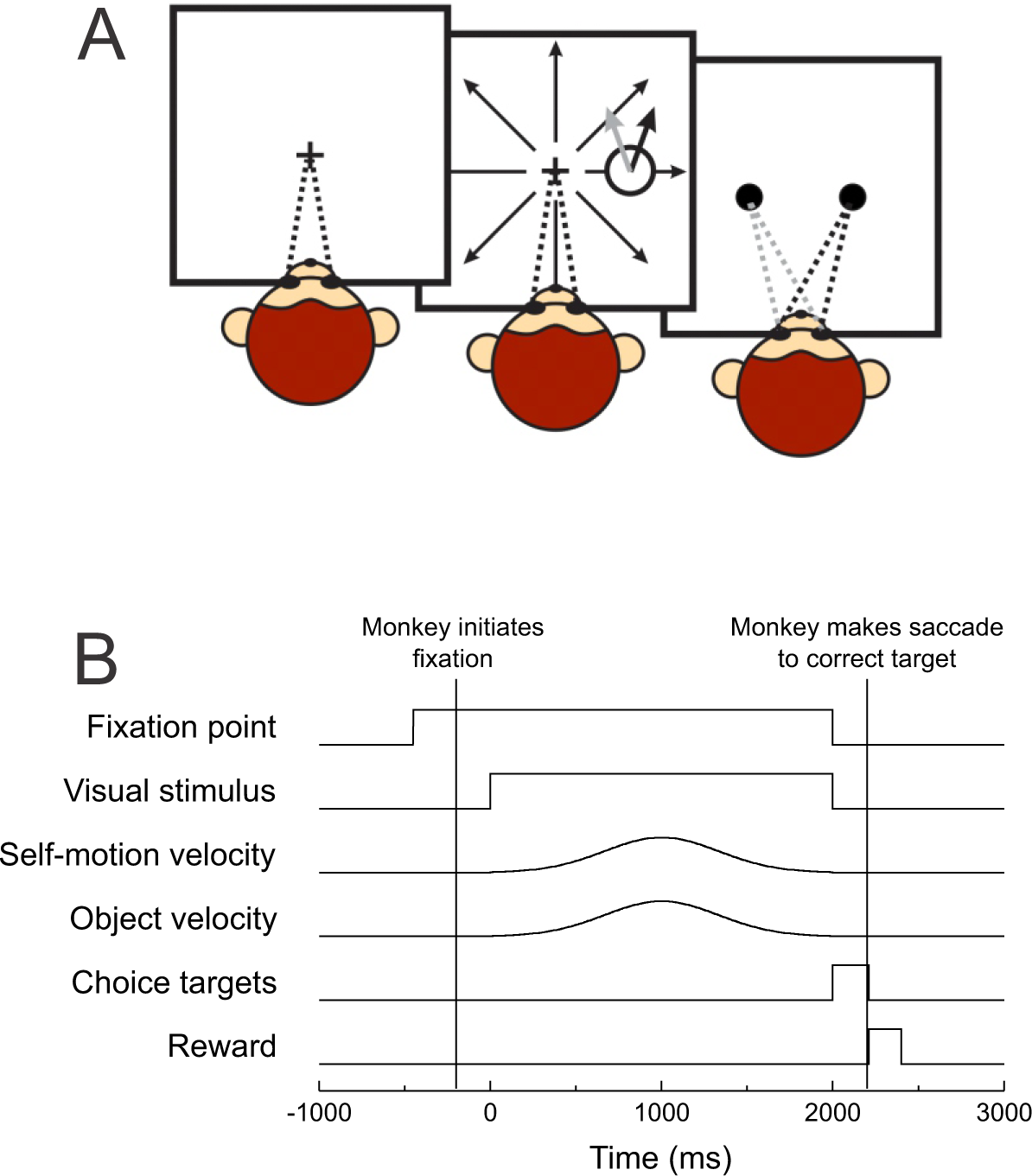
**Schematic illustration of the object direction discrimination task.** (A) Each trial initiated when a fixation target appeared and the monkey fixated on the target. The monkey was required to maintain fixation during the presentation of a stimulus, which consisted of an object moving upward obliquely and a global optic flow field simulating forward or backward self-motion. At the end of the stimulus presentation, two choice targets appeared. The monkey was required to make a saccade to one of the targets indicating whether the object’s motion was rightward or leftward of vertical. (B) Timeline of events within each trial.

**Supplementary Figure 3.**
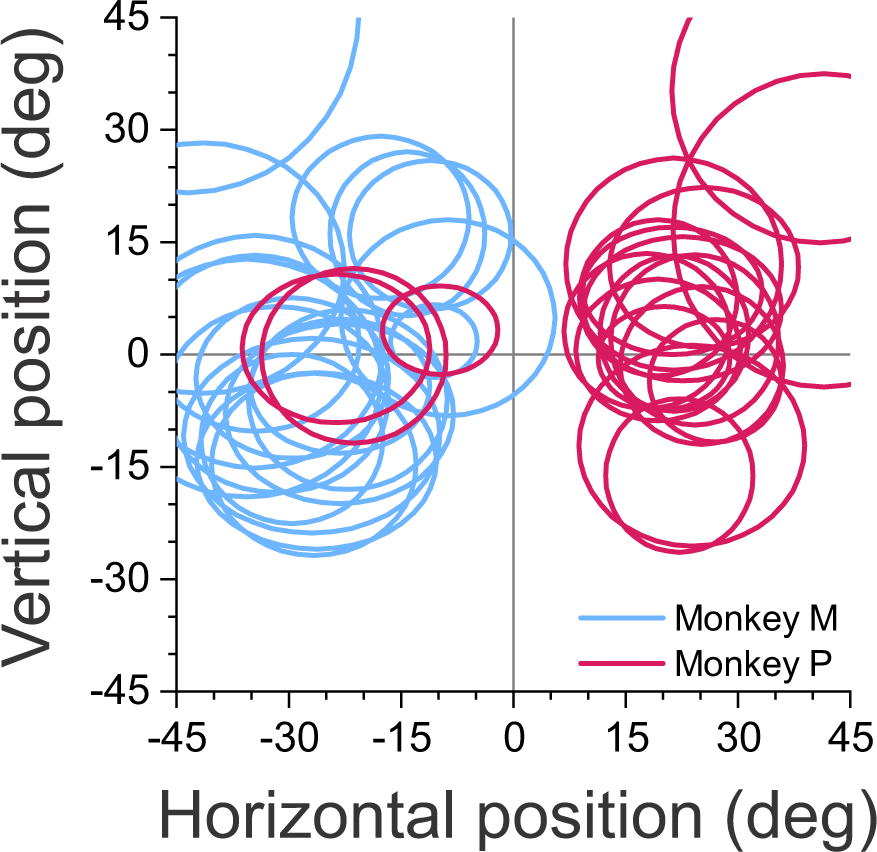
**Receptive field locations for each experimental session.** Each ellipse represents the average receptive field location and size across all units recorded within a session, computed from the average parameters of 2-dimensional (2D) Gaussian fits. Each ellipse corresponds to a cross-section through the average 2D Gaussian function at the half-maximal response amplitude. Coordinate (0, 0) represents the center of the screen and the location of the fixation target. Color indicates monkey identity; teal: monkey M, purple: monkey P.

**Supplementary Figure 4.**
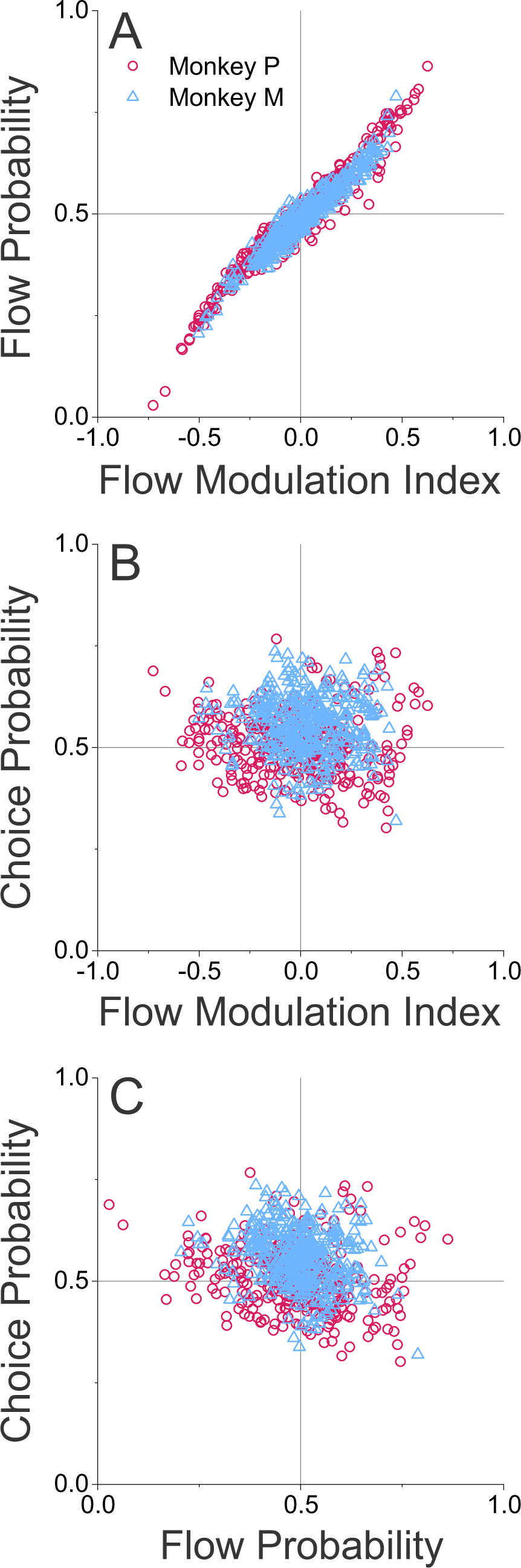
**Relationships between FMI, Flow Probability, and Choice Probability.** In all panels, symbol color and shape denote monkeys M (teal triangles) and P (purple circles). (A) Flow probability (FP) is highly correlated with flow modulation index (FMI) for both monkey P (Pearson correlation, R = 0.97, P = 3.2×10^-205^, N=315) and monkey M (R = 0.96, P = 1.2×10^-241^, N=410). (B) Choice probability (CP) is poorly correlated with FMI for both monkey P (Pearson correlation, R = -0.08, P = 0.147, N=315) and monkey M (R = -0.07, P = 0.176, N=410). (C) CP is weakly negatively correlated with FP for both monkey P (Pearson correlation, R = -0.22, P = 5.0×10^-5^, N=315) and monkey M (R = -0.25, P = 1.5×10^-7^, N=410).

**Supplementary Figure 5.**
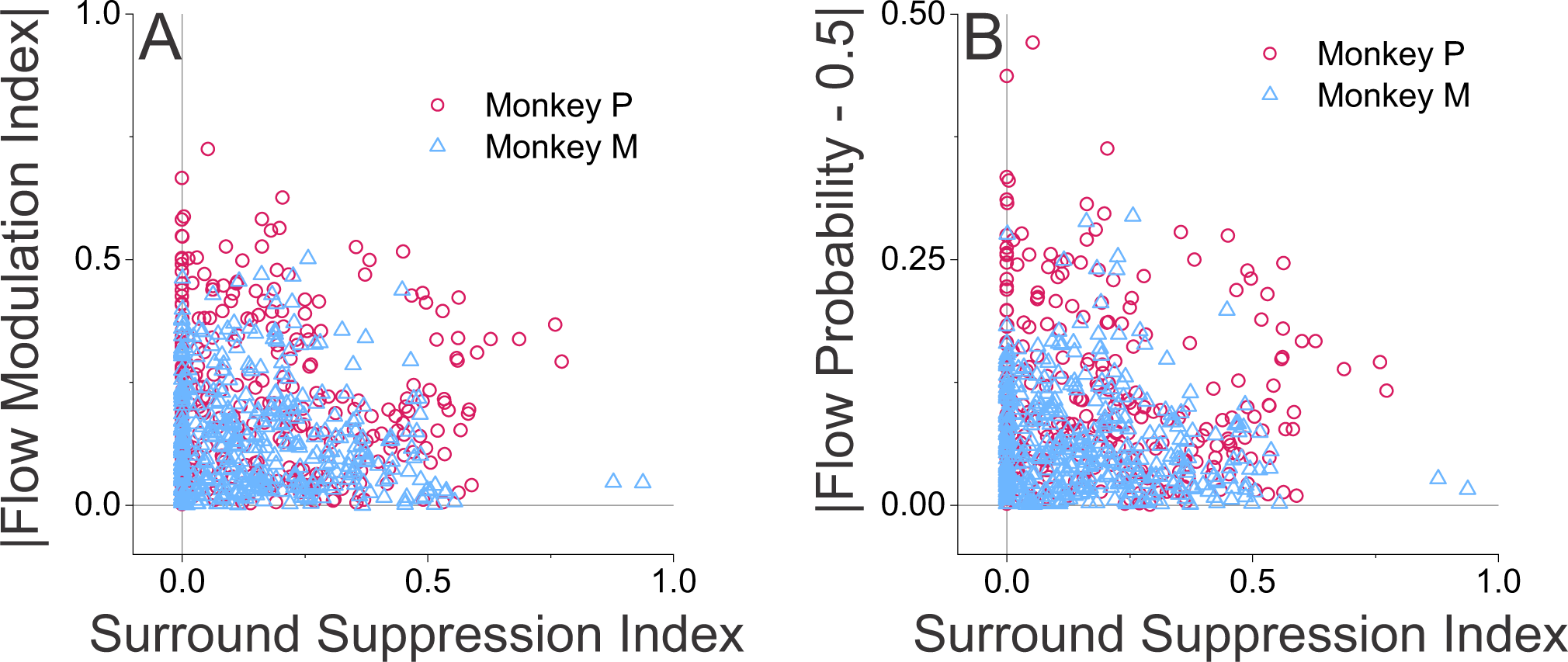
Optic flow modulation in MT is not correlated with surround suppression. (A) Absolute value of FMI is plotted as a function of surround suppression index. Symbol color and shape denote monkeys M (teal triangles) and P (purple circles). (B) The absolute value of flow probability, after subtracting 0.5, is plotted against surround suppression index (format as in panel B). If effects of background optic flow on MT responses were explained by surround suppression, we would expect to see positive correlations in these plots.

**Movie 1. Examples of visual stimuli.** This video shows examples of visual stimuli from the main flow-parsing experiment. A sequence of 4 trials is shown. In each trial, the target object patch (composed of small yellow triangles) moves vertically. Background optic flow (shown as a red/green anaglyph for stereo viewing) simulates self-motion that alternates between forward (expanding optic flow) and backward (contracting optic flow). The small yellow cross at the center of the display is the target to be fixated while viewing the stimuli. The prediction of flow parsing is that the target object should appear to be moving up-left during forward self-motion (first and third trials) and up-right during backward self-motion (second and fourth trials).

## REFERENCES

Albright, T.D., and Desimone, R. (1987). Local precision of visuotopic organization in the middle temporal area (MT) of the macaque. Experimental Brain Research 65, 582–592.

Albright, T.D., Desimone, R., and Gross, C.G. (1984). Columnar organization of directionally selective cells in visual area MT of the macaque. Journal of Neurophysiology 51, 16–31. 10.1152/jn.1984.51.1.16.

Allman, J., Miezin, F., and McGuinness, E. (1985). Direction- and velocity-specific responses from beyond the classical receptive field in the middle temporal visual area (MT). Perception 14, 105–126.

Born, R.T. (2000). Center-surround interactions in the middle temporal visual area of the owl monkey. Journal of Neurophysiology 84, 2658–2669. 10.1152/jn.2000.84.5.2658.

Bradley, D.C., and Andersen, R.A. (1998). Center-surround antagonism based on disparity in primate area MT. Journal of Neuroscience 18, 7552–7565.

Bradley, D.C., Maxwell, M., Andersen, R.A., Banks, M.S., and Shenoy, K.V. (1996). Mechanisms of heading perception in primate visual cortex. Science 273, 1544–1547.

Bremmer, F., Duhamel, J.R., Ben Hamed, S., and Graf, W. (2002). Heading encoding in the macaque ventral intraparietal area (VIP). European Journal of Neuroscience 16, 1554–1568.

Britten, K.H. (2008). Mechanisms of self-motion perception. Annual Review of Neuroscience 31, 389–410. 10.1146/annurev.neuro.29.051605.112953.

Britten, K.H., Newsome, W.T., Shadlen, M.N., Celebrini, S., and Movshon, J.A. (1996). A relationship between behavioral choice and the visual responses of neurons in macaque MT. Visual Neuroscience 13, 87–100.

Britten, K.H., and Van Wezel, R.J. (2002). Area MST and heading perception in macaque monkeys. Cerebral Cortex 12, 692–701.

Celebrini, S., and Newsome, W.T. (1995). Microstimulation of extrastriate area MST influences performance on a direction discrimination task. Journal of Neurophysiology 73, 437–448.

Chen, A., DeAngelis, G.C., and Angelaki, D.E. (2011). Representation of vestibular and visual cues to self-motion in ventral intraparietal cortex. Journal of Neuroscience 31, 12036–12052. 10.1523/JNEUROSCI.0395-11.2011.

Chen, A., Deangelis, G.C., and Angelaki, D.E. (2013). Functional specializations of the ventral intraparietal area for multisensory heading discrimination. J Neurosci 33, 3567–3581. 10.1523/JNEUROSCI.4522-12.2013.

DeAngelis, G.C., and Newsome, W.T. (1999). Organization of disparity-selective neurons in macaque area MT. Journal of Neuroscience 19, 1398–1415.

DeAngelis, G.C., and Uka, T. (2003). Coding of horizontal disparity and velocity by MT neurons in the alert macaque. Journal of Neurophysiology 89, 1094–1111.10.1152/jn.00717.2002.

Dokka, K., MacNeilage, P.R., DeAngelis, G.C., and Angelaki, D.E. (2015). Multisensory self-motion compensation during object trajectory judgments. Cereb Cortex 25, 619–630. 10.1093/cercor/bht247.

Duffy, C.J., and Wurtz, R.H. (1991). Sensitivity of MST neurons to optic flow stimuli. I. A continuum of response selectivity to large-field stimuli. Journal of Neurophysiology 65, 1329–1345.

Duffy, C.J., and Wurtz, R.H. (1995). Responses of monkey MST neurons to optic flow stimuli with shifted centers of motion. Journal of Neuroscience 15, 5192–5208.

Fajen, B.R., Parade, M.S., and Matthis, J.S. (2013). Humans perceive object motion in world coordinates during obstacle avoidance. Journal of Vision 13. 10.1167/13.8.25.

Fattori, P., Pitzalis, S., and Galletti, C. (2009). The cortical visual area V6 in macaque and human brains. J Physiol Paris 103, 88–97. 10.1016/j.jphysparis.2009.05.012.

Fetsch, C.R., Pouget, A., DeAngelis, G.C., and Angelaki, D.E. (2012). Neural correlates of reliability-based cue weighting during multisensory integration. Nature Neuroscience 15, 146–154. 10.1038/nn.2983.

Field, D.T., Biagi, N., and Inman, L.A. (2020). The role of the ventral intraparietal area (VIP/pVIP) in the perception of object-motion and self-motion. Neuroimage 213, 116679. 10.1016/j.neuroimage.2020.116679.

Foulkes, A.J., Rushton, S.K., and Warren, P.A. (2013). Flow parsing and heading perception show similar dependence on quality and quantity of optic flow. Frontiers in Behavioral Neuroscience 7. 10.3389/fnbeh.2013.00049.

Galletti, C., Battaglini, P.P., and Fattori, P. (1995). Eye position influence on the parieto-occipital area PO (V6) of the macaque monkey. European Journal of Neuroscience 7, 2486–2501. 10.1111/j.1460-9568.1995.tb01047.x.

Gibson, J.J. (1950). The Perception of the Visual World (Houghton Mifflin).

Gu, Y., Angelaki, D.E., and DeAngelis, G.C. (2008). Neural correlates of multisensory cue integration in macaque MSTd. Nature Neuroscience 11, 1201–1210. 10.1038/nn.2191.

Gu, Y., DeAngelis, G.C., and Angelaki, D.E. (2012). Causal links between dorsal medial superior temporal area neurons and multisensory heading perception. Journal of Neuroscience 32, 2299–2313. 10.1523/Jneurosci.5154-11.2012.

Gu, Y., Watkins, P.V., Angelaki, D.E., and DeAngelis, G.C. (2006). Visual and nonvisual contributions to three-dimensional heading selectivity in the medial superior temporal area. Journal of Neuroscience 26, 73–85. 10.1523/JNEUROSCI.2356-05.2006.

Hatsopoulos, N.G., and Warren, W.H. (1991). Visual navigation with a neural network. Neural Networks 4, 303–317. 10.1016/0893-6080(91)90068-G.

Kim, H.R., Angelaki, D.E., and DeAngelis, G.C. (2015). A novel role for visual perspective cues in the neural computation of depth. Nat Neurosci 18, 129–137. 10.1038/nn.3889.

Kim, H.R., Angelaki, D.E., and DeAngelis, G.C. (2017). Gain modulation as a mechanism for coding depth from motion parallax in macaque area MT. Journal of Neuroscience 37, 8180–8197. 10.1523/JNEUROSCI.0393-17.2017.

Kohn, A., and Movshon, J.A. (2003). Neuronal adaptation to visual motion in area MT of the macaque. Neuron 39, 681–691. 10.1016/s0896-6273(03)00438-0.

Kourtzi, Z., and Kanwisher, N. (2000). Activation in human MT/MST by static images with implied motion. Journal of Cognitive Neuroscience 12, 48–55. 10.1162/08989290051137594.

Kozhemiako, N., Nunes, A.S., Samal, A., Rana, K.D., Calabro, F.J., Hämäläinen, M.S., Khan, S., and Vaina, L.M. (2020). Neural activity underlying the detection of an object movement by an observer during forward self-motion: Dynamic decoding and temporal evolution of directional cortical connectivity. Prog Neurobiol 195, 101824. 10.1016/j.pneurobio.2020.101824.

Krekelberg, B., Dannenberg, S., Hoffmann, K.P., Bremmer, F., and Ross, J. (2003). Neural correlates of implied motion. Nature 424, 674–677. 10.1038/nature01852.

Layton, O.W., and Browning, N.A. (2014). A unified model of heading and path perception in primate MSTd. PLoS Computational Biology 10, 1–20. 10.1371/journal.pcbi.1003476.

Layton, O.W., and Fajen, B.R. (2016a). Competitive Dynamics in MSTd: A Mechanism for Robust Heading Perception Based on Optic Flow. PLoS Computational Biology 12, 1–37. 10.1371/journal.pcbi.1004942.

Layton, O.W., and Fajen, B.R. (2016b). A neural model of MST and MT explains perceived object motion during self-motion. Journal of Neuroscience 36, 8093–8102. 10.1523/JNEUROSCI.4593-15.2016.

Layton, O.W., Mingolla, E., and Browning, N.A. (2012). A motion pooling model of visually guided navigation explains human behavior in the presence of independently moving objects. Journal of Vision 12, 1–19. 10.1167/12.1.20.

Layton, O.W., and Niehorster, D.C. (2019). A model of how depth facilitates scene-relative object motion perception. Plos Computational Biology 15. 10.1371/journal.pcbi.1007397.

Longuet-Higgins, H.C., and Prazdny, K. (1980). The interpretation of a moving retinal image. Proceedings of the Royal Society of London. Series B, Biological Sciences 208, 385–397.

Luo, J., He, K., Andolina, I.M., Li, X., Yin, J., Chen, Z., Gu, Y., and Wang, W. (2019). Going with the flow: the neural mechanisms underlying illusions of complex-flow motion. Journal of Neuroscience 39, 2664–2685. 10.1523/JNEUROSCI.2112-18.2019.

Matsumiya, K., and Ando, H. (2009). World-centered perception of 3D object motion during visually guided self-motion. Journal of Vision 9, 15 11–13. 10.1167/9.1.15.

Maunsell, J.H.R., and Van Essen, D.C. (1983a). The connections of the middle temporal visual area (MT) and their relationship to a cortical hierarchy in the macaque monkey. Journal of Neuroscience 3, 2563–2586.

Maunsell, J.H.R., and Van Essen, D.C. (1983b). Functional properties of neurons in middle temporal visual area of the macaque monkey. I. Selectivity for stimulus direction, speed, and orientation. Journal of Neurophysiology 49, 1127–1147.

Niehorster, D.C., and Li, L. (2017). Accuracy and tuning of flow parsing for visual perception of object motion during self-motion. i-Perception 8. 10.1177/2041669517708206.

Nienborg, H., Cohen, M.R., and Cumming, B.G. (2012). Decision-related activity in sensory neurons: Correlations among neurons and with behavior. Annual Reviews of Neuroscience 35, 463–483. 10.1146/annurev-neuro-062111-150403.

Nogueira, R., Abolafia, J.M., Drugowitsch, J., Balaguer-Ballester, E., Sanchez-Vives, M.V., and Moreno-Bote, R. (2017). Lateral orbitofrontal cortex anticipates choices and integrates prior with current information. Nat Commun 8, 14823. 10.1038/ncomms14823.

Nover, H., Anderson, C.H., and DeAngelis, G.C. (2005). A logarithmic, scale-invariant representation of speed in macaque middle temporal area accounts for speed discrimination performance. Journal of Neuroscience 25, 10049–10060. 10.1523/JNEUROSCI.1661-05.2005.

Peltier, N.E., Angelaki, D.E., and DeAngelis, G.C. (2020). Optic flow parsing in the macaque monkey. Journal of Vision 20, 1–27. 10.1167/jov.20.10.8.

Perrone, J.A. (1992). Model for the computation of self-motion in biological systems. J Opt Soc Am A 9, 177–194.

Perrone, J.A. (2012). A neural-based code for computing image velocity from small sets of middle temporal (MT/V5) neuron inputs. Journal of Vision 12, 1–31. 10.1167/12.8.1.

Perrone, J.A., and Stone, L.S. (1994). A model for self-motion estimation within primate extrastriate visual cortex. Vision Research 34, 2917–2938.

Perrone, J.A., and Stone, L.S. (1998). Emulating the visual receptive-field properties of MST neurons with a template model of heading estimation. Journal of Neuroscience 18, 5958–5975.

Pitzalis, S., Serra, C., Sulpizio, V., Committeri, G., de Pasquale, F., Fattori, P., Galletti, C., Sepe, R., and Galati, G. (2020). Neural bases of self- and object-motion in a naturalistic vision. Hum Brain Mapp 41, 1084–1111. 10.1002/hbm.24862.

Prince, S.J.D., Pointon, A.D., Cumming, B.G., and Parker, A.J. (2002). Quantitative analysis of the responses of V1 neurons to horizontal disparity in dynamic random-dot stereograms. Journal of Neurophysiology 87, 191–208. 10.1152/jn.00465.2000.

Purushothaman, G., and Bradley, D.C. (2005). Neural population code for fine perceptual decisions in area MT. Nature Neuroscience 8, 99–106. 10.1038/nn1373.

Rodman, H.R., and Albright, T.D. (1989). Single-unit analysis of pattern-motion selective properties in the middle temporal visual area (MT). Experimental Brain Research 75, 53–64. 10.1007/bf00248530.

Rogers, C., Rushton, S.K., and Warren, P.A. (2017). Peripheral visual cues contribute to the perception of object movement during self-movement. i-Perception 8. 10.1177/2041669517736072.

Royden, C.S. (1997). Mathematical analysis of motion-opponent mechanisms used in the determination of heading and depth. Journal of the Optical Society of America A 14, 2128–2143. 10.1364/josaa.14.002128.

Royden, C.S., and Holloway, M.A. (2014). Detecting moving objects in an optic flow field using direction- and speed-tuned operators. Vision Research 98, 14–25.

Rushton, S.K., and Warren, P.A. (2005). Moving observers, relative retinal motion and the detection of object movement. Current Biology 15, R542–543. 10.1016/j.cub.2005.07.020.

Sasaki, R., Anzai, A., Angelaki, D.E., and DeAngelis, G.C. (2020). Flexible coding of object motion in multiple reference frames by parietal cortex neurons. Nature Neuroscience. 10.1038/s41593-020-0656-0.

Schlack, A., and Albright, T.D. (2007). Remembering visual motion: neural correlates of associative plasticity and motion recall in cortical area MT. Neuron 53, 881–890. 10.1016/j.neuron.2007.02.028.

Shivkumar, S., DeAngelis, G.C., and Haefner, R.M. (2023). Hierarchical motion perception as causal inference. Under review.

Stoner, G.R., and Albright, T.D. (1992). Neural correlates of perceptual motion coherence. Nature 358, 412–414. 10.1038/358412a0.

Uka, T., and DeAngelis, G.C. (2004). Contribution of area MT to stereoscopic depth perception: choice-related response modulations reflect task strategy. Neuron 42, 297–310. 10.1016/s0896-6273(04)00186-2.

Van den Berg, A.V. (1992). Robustness of perception of heading from optic flow. Vision Research 32, 1285–1296. 10.1016/0042-6989(92)90223-6.

Van Essen, D.C., Drury, H.A., Dickson, J., Harwell, J., Hanlon, D., and Anderson, C.H. (2001). An integrated software suite for surface-based analyses of cerebral cortex. J Am Med Inform Assoc 8, 443–459.

Van Essen, D.C., Maunsell, J.H., and Bixby, J.L. (1981). The middle temporal visual area in the macaque: myeloarchitecture, connections, functional properties and topographic organization. J Comp Neurol 199, 293–326. 10.1002/cne.901990302.

Van Wezel, R.J., and Britten, K.H. (2002). Motion adaptation in area MT. Journal of Neurophysiology 88, 3469–3476. 10.1152/jn.00276.2002.

Warren, P.A., and Rushton, S.K. (2007). Perception of object trajectory: Parsing retinal motion into self and object movement components. Journal of Vision 7. 10.1167/7.11.2.

Warren, P.A., and Rushton, S.K. (2009a). Optic flow processing for the assessment of object movement during ego movement. Current Biology 19, 1555–1560. 10.1016/j.cub.2009.07.057.

Warren, P.A., and Rushton, S.K. (2009b). Perception of scene-relative object movement: Optic flow parsing and the contribution of monocular depth cues. Vision Research 49, 1406–1419. 10.1016/j.visres.2009.01.016.

Warren, W.H., Blackwell, A.W., Kurtz, K.J., Hatsopoulos, N.G., and Kalish, M.L. (1991). On the sufficiency of the velocity field for perception of heading. Biological Cybernetics 65, 311–320.

Zaidel, A., DeAngelis, G.C., and Angelaki, D.E. (2017). Decoupled choice-driven and stimulus-related activity in parietal neurons may be misrepresented by choice probabilities. Nat Commun 8, 715. 10.1038/s41467-017-00766-3.

Zhang, T., and Britten, K.H. (2004). Clustering of selectivity for optic flow in the ventral intraparietal area. Neuroreport 15, 1941–1945. 10.1097/00001756-200408260-00022.

Zhang, T., and Britten, K.H. (2010). The responses of VIP neurons are sufficiently sensitive to support heading judgments. Journal of Neurophysiology 103, 1865–1873. 10.1152/jn.00401.2009.

